# A tunable aqueous architecture modulates functional output in biomolecular condensates

**DOI:** 10.64898/2026.05.13.724666

**Authors:** Moeka Sasazawa, Mechi Chen, Rui Zeng, Uvarov Denis, Sujata Bais, Jillian Hoffstadt, Julian von Hofe, Norah Hoffmann, Yulia Volkova, Saumya Saurabh

## Abstract

Biomolecular condensates organize cellular biochemistry, yet the principles governing their internal solvent architectures remain poorly understood. Most current models focus on macromolecular scaffolds while treating the solvent as a passive, spatially uniform background. Here, we introduce Condensate Spatial Topography via Emission Lifetimes (ConSTEL) to map the continuous solvent polarity landscape inside biomolecular condensates. Using PopZ as a model system, we show that the condensate interior contains a persistent, tunable mosaic of aqueous environments whose apparent polarity, reported by Nile Red fluorescence lifetimes, is organized by thermodynamic state and chemical cues. This microphase-separated solvent architecture defines distinct mesoscale rheological regimes, with intermediate aqueous niches supporting fast, confined tracer motion and highly polar or non-polar extremes forming a slower, viscoelastic mesh. We further demonstrate that drug-like small molecules partition non-uniformly across this landscape according to their physicochemical properties, and that exceeding local solubility limits drives “reciprocal sculpting”, in which mismatched guests remodel the host solvent architecture. Together, these results highlight internal solvent organization as an active, tunable determinant of condensate material properties, molecular transport, and partitioning, and suggest that predictive models of condensate function and pharmacology would benefit from incorporating the spatial arrangement of solvent environments alongside bulk composition.

## Introduction

Biomolecular condensates have emerged as a primary frontier in targeted pharmacology due to their capacity to selectively compartmentalize and concentrate small-molecule therapeutics [1–3]. While bulk partition coefficients (*C_in_/C_out_*) provide a highly useful metric for quantifying the overall thermodynamic recruitment of small molecules [4–6], these mean-field descriptors inherently average over the entire droplet volume. By construction, they obscure any internal spatial structure and implicitly treat the condensate interior as a uniformly shared hydrophobic phase [7–9]. As a result, current predictive models relying exclusively on bulk *C_in_/C_out_* face a critical limitation: they cannot distinguish between chemically distinct niches within the same condensate, nor can they explain how local environments shape mesoscale transport and pharmacological access.

Recent theoretical, bulk spectroscopy, and single-molecule microscopy studies have firmly established that biomolecular condensates are profoundly heterogeneous across multiple spatiotemporal scales [10–15]. However, this emerging picture is still fundamentally macromolecule-centric: it focuses on how scaffolds and clients phase separate, form networks, and create domains, while relegating the interpenetrating aqueous phase to a largely passive role. In this view, the solvent is often treated as a uniform background whose only function is to support macromolecular interactions, leaving its own structure and dynamics effectively uncharacterized.

At the same time, a complementary body of work makes it clear that the solvent is neither uniform nor passive inside condensates. Terahertz (THz) spectroscopy and atomistic simulations demonstrate that structured water actively drives phase separation, while water retained in the direct protein hydration layer is dynamically retarded compared to the bulk [16–18]. Similarly, ultrafast two-dimensional infrared (2D IR) spectroscopy shows that intra-condensate water is partitioned into distinct dynamic populations, implying confinement within nanometer-scale pores [19]. Furthermore, hyperspectral mapping has quantified a macroscopic reduction in water dipolar relaxation and effective dielectric permittivity within condensates [20, 21].

While these foundational studies confirm that the scaffold and water are organized differently inside the dense phase, a critical observational gap persists. Single-molecule techniques resolve the complex spatial architecture of the macromolecular scaffold but remain chemically blind to the solvent. Conversely, current solvent-resolving methods successfully capture the altered dynamics of the internal water, but they either average these dynamics across the entire droplet or report uniform macroscopic environments [19, 21]. Because of this disconnect between spatial and chemical resolution, the field has only been able to indirectly infer the coexistence of distinct aqueous sub-populations. Consequently, the continuous topology of this internal scaffold-solvent network, which is the landscape that molecules experience within the condensate, remains essentially unresolved.

Here, we introduce Condensate Spatial Topography via Emission Lifetimes (Con-STEL), a time-averaged fluorescence lifetime imaging (FLIM) method that directly visualizes the continuous internal macromolecule-solvent architecture of biomolecular condensates. ConSTEL leverages the solvatochromic dye Nile Red, whose excited-state lifetime is exquisitely sensitive to solvent-mediated hydrogen-bond quenching and can be calibrated to an *E_T_* (30)-like scale as a spatially resolved proxy for local water accessibility and polarity [22–24]. Applying ConSTEL to condensates formed by the bacterial hub protein PopZ [25, 26], we reveal a persistent, tunable mosaic of aqueous microenvironments organized by thermodynamic state and chemical cues. By correlating this solvent topology with nanoparticle transport and small-molecule distribution, we show that internal solvent organization is a central design variable linking condensate thermodynamics, mesoscale rheology, and molecular partitioning. Moreover, when local solubility limits are exceeded, mismatched guest small-molecules can, in turn, remodel this solvent architecture.

## Results

### Time-averaged FLIM reveals a stable solvent architecture governed by hydrogen-bonding in biomolecular condensates

To experimentally map the continuous chemical environments in biomolecular condensates, we used PopZ as a model system with established structural and dynamic heterogeneity [27–29]. Single-molecule localization and image correlation spectroscopy reveal pronounced spatial and temporal heterogeneities within PopZ condensates [14], while structural studies have resolved an underlying filamentous ultrastructure [30, 31].

We first visualized the system using standard confocal fluorescence microscopy of the solvatochromic dye Nile Red.[32] Although intensity maps display localized dye accumulation and spatial heterogeneity (Fig. 1a), this readout is inherently ambiguous because intensity alone conflates changes in local probe concentration, aggregation state, and excitation/detection conditions with genuine differences in solvent chemistry. We therefore turned to time-averaged fluorescence lifetime mapping, which is largely independent of probe concentration and provides a more direct, quantitative readout of the local chemical environment.

**Fig. 1:**
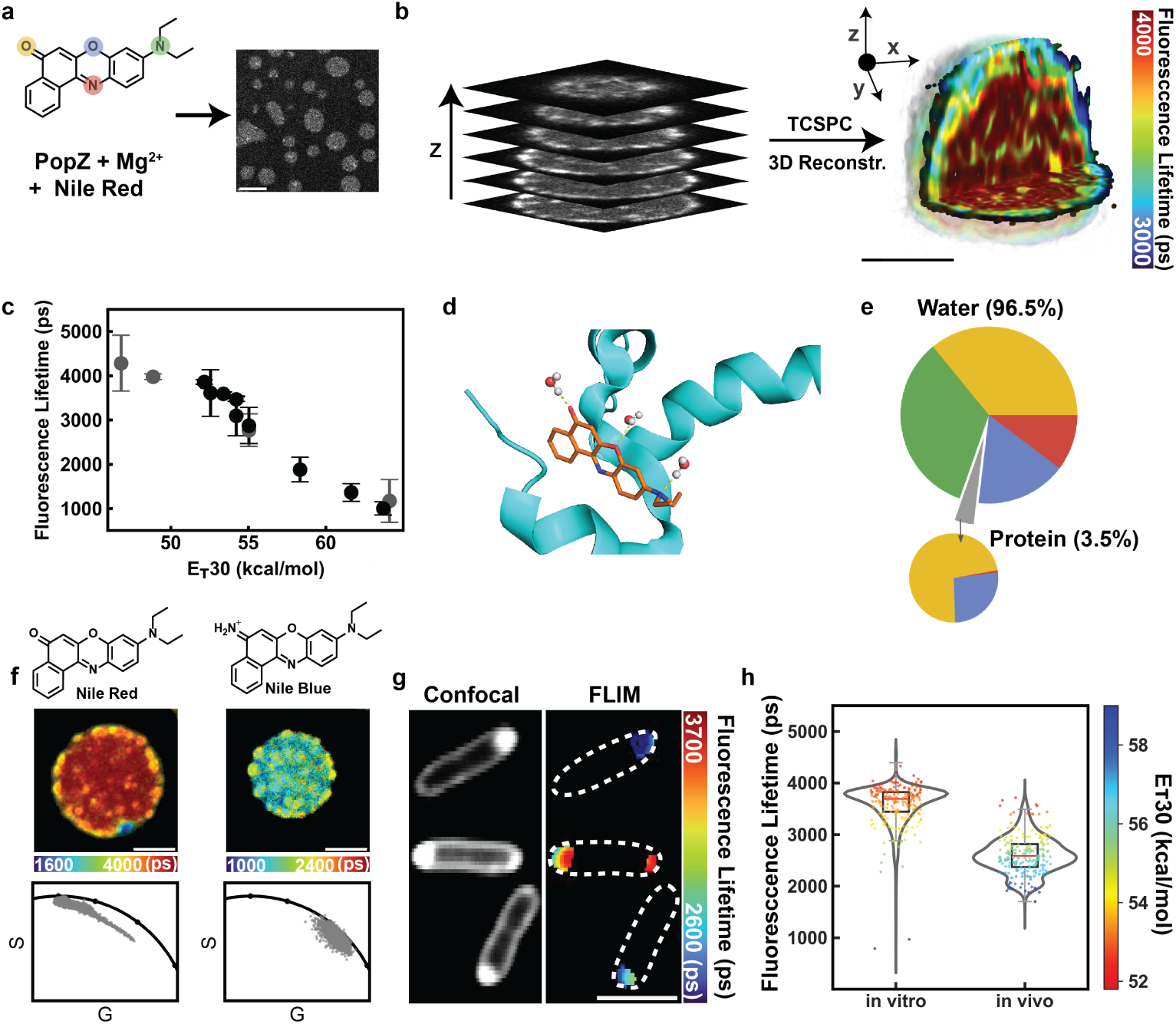
Time-averaged FLIM maps complex protein–solvent architecture in PopZ condensates. **a,** Schematic of ConSTEL. Nile Red is mixed with PopZ and Mg^2+^ to form biomolecular condensates for confocal fluorescence microscopy (Scale bar, 5 µm). **b,** Volumetric lifetime map of a single PopZ condensate generated via TCSPC-acquired *z*-stacks followed by deconvolution. **c,** Calibration curve showing the relationship between Nile Red fluorescence lifetime and the solvent polarity parameter *E_T_* (30) across a range of solvents. Data are mean ± standard deviation (*n >* 10,000 pixels per point; representative of two independent experiments). **d,** All-atom molecular dynamics (MD) simulation snapshot of Nile Red located within a PopZ dense phase, illustrating a local solvation shell in which water molecules preferentially coordinate the fluorophore. **e,** MD-derived hydrogen-bonding hierarchy. Water molecules account for 96.5% of total hydrogen bonds to Nile Red; protein contacts contribute 3.5%. Colors match specific atoms in Nile Red (panel **a**). **f,** Comparative imaging of PopZ condensates using Nile Red (left) and Nile Blue (right), with corresponding chemical structures (top), lifetime maps (middle), and phasor plots (bottom). Nile Red provides a broader dynamic range of lifetimes than Nile Blue. **g,** Confocal and FLIM images of Nile Red-labeled PopZ condensates in live *E. coli*, demonstrating conservation of heterogeneous architecture *in vivo*. Membrane-associated signal of Nile Red is masked. **h,** Violin plots of pixel lifetime distributions in *in vitro* and *in vivo* PopZ condensates, highlighting broad, overlapping polarity ranges (*n* = 684,489 pixels (*in vivo*) and *n* = 773,499 pixels (*in vitro*) from three independent measurements, Scale bars, 2 µm (**e–g**)).

Strikingly, time-averaged 3D *z*-stack reconstructions revealed a spatially stable mosaic of discrete lifetimes on the order of 3000–4000 ps within single PopZ condensates (Fig. 1b, Supplementary video 1). To relate these signals to solvent polarity, we rigorously calibrated the fluorescence lifetime of Nile Red against the empirical *E_T_* (30) polarity scale [33] via pure-solvent titrations (Fig. 1c; Supplementary Fig. S1a–e). While Nile Red photophysics have been interpreted through free-volume–dependent twisted or planar Intramolecular Charge-Transfer (ICT) models to probe local viscosity [34, 35], our pure-solvent calibrations and time-dependent density functional theory (TDDFT) calculations do not support such a mechanism (Supplementary Note 1). Across a broad range of solvents, the fluorescence lifetime shows only modest variation with dielectric constant, viscosity, and H-bond acceptor content, but decreases sharply with increasing hydrogen-bond donor density [22]. To understand the role of hydrogen bonding in dense-phase-like environments, we performed all-atom molecular dynamics (MD) simulations of Nile Red and PopZ at experimental dense phase concentrations [14]. These simulations reveal that 96.5% of transient hydrogen bonds involving the probe originate from interfacial water molecules, with only a minor contribution from PopZ side chains (Fig. 1d,e; Supplementary Video 2, Supplementary Fig. S1f–l). This simulation-based picture is consistent with Nile Red primarily reporting on the aqueous phase rather than protein-mediated interactions in the dense condensate. Consequently, ConSTEL provides a direct, quantitative readout of nanoscale water accessibility.

These lifetime features are conserved over extended continuous acquisition windows, indicating that the protein-solvent architecture is structurally persistent on experimental timescales (Supplementary Fig. S2a). At the same time, fluorescence recovery after photobleaching (FRAP) shows that Nile Red exhibits similar mobility across environments of varying polarity (Supplementary Fig. S2b). These observations indicate that while molecular constituents exchange rapidly, the underlying pattern of solvent polarities is time-averaged and stable, behaving as a quasi-static landscape through which the condensate constituents move.

To test whether ConSTEL perturbs the condensates or introduces systematic lifetime biases, we combined label-free holography, continuous photon tracking, and Monte Carlo FLIM simulations (Supplementary Note 2). Continuous photon tracking indicates that Nile Red behaves as a freely exchanging reporter, showing a cumulative increase in signal over time (Supplementary Fig. S2c). Label-free holography confirmed that Nile Red does not measurably perturb the native phase equilibrium (Supplementary Fig. S2d). Following standard FLIM practice [36], we sampled each condensate with pixels smaller than the diffraction-limited point-spread function (50 nm pixels) and observed stable features between 100-300 nm (Supplementary Fig. S2e). We used Monte Carlo simulations to quantify the impact of this spatial oversampling on life-time precision. In the photon-count regime of our experiments (*>*100 photons at the peak of the photon-count histogram per pixel)), these simulations showed that life-time estimates from the super-sampled image remain accurate to within ∼5% at the length scale of the solvent microdomains (Supplementary Fig. S2f–h), indicating that the observed spatial complexity reflects genuine physical heterogeneity rather than photophysical, optical or statistical artifacts.

To isolate the role of aromatic interactions with the probe on the the lifetime map, we replaced Nile Red with Nile Blue. Nile Blue shares the same aromatic core as Nile Red, and can therefore engage similar *π*–*π* and cation–*π* interactions, but lacks lifetime sensitivity to hydrogen-bond–mediated quenching. Nile Blue exhibited a substantially narrowed distribution of lifetimes and much weaker structural contrast than Nile Red (Fig. 1f), indicating that the pronounced lifetime mosaic primarily reports the hydrogen-bonded water network rather than transient aromatic stacking.

Functionally, this spatially partitioned, solvent-rich architecture can locally influence chemical reactivity. The localized polar niches correlate with the water-dependent hydrolysis of fluorescein diacetate (Supplementary Fig. S2i). Importantly, this heterogeneous solvent architecture is conserved within PopZ condensates in live *E. coli* (Fig. 1g,h). Taken together, these data establish that PopZ condensates contain a spatially stable, chemically structured solvent phase that is preserved *in vivo* and can locally modulate aqueous reactivity. Having defined this archetypal solvent mosaic in PopZ, we next asked whether similar architectures arise in other condensate systems and how thermodynamic control knobs tune their topology.

### Solvent architecture as a tunable thermodynamic state in condensates

Applying ConSTEL beyond PopZ, we first asked whether structured aqueous architectures are unique to this bacterial scaffold or represent a general feature of condensates. A complex coacervate (polyA–spermine) [37] and a folded-protein condensate (BSA–PEG)[38], both displayed spatially stable mosaics of solvent polarities that could be segmented into multiple *E_T_* (30) bins (Supplementary Fig. S2j). Thus, coexisting solvent microenvironments appear to be a common feature of these phase-separated network fluids. At the same time, the quantitative *E_T_* (30) ranges and bin occupancies depend on the physicochemical properties of the underlying scaffolds and co-solutes, with PopZ–MgCl_2_, BSA–PEG, and polyA–spermine condensates each showing distinct polarity boundaries. These observations are consistent with different condensates having system-specific solvent landscapes.

Focusing on PopZ, we tested whether its solvent topology acts as a thermodynamic state variable determined by the system’s position in phase space. We mapped the internal solvent architecture of PopZ condensates across a range of saturation conditions by systematically varying PopZ and Mg^2+^ concentrations (Fig. 2a). To quantify the resulting topological changes, we first analyzed the global frequency distribution of *E_T_* (30) microenvironments to define polarity thresholds (Supplementary Fig. S3a). We then segmented the continuous *E_T_* (30) landscape into four polarity bins—from Bin 1 (most hydrophobic) to Bin 4 (most aqueous)—and computed the fractal dimension *D_f_*of each bin to capture its spatial connectivity (Fig. 2b; Supplementary Fig. S3b). Condensates near the phase boundary (binodal) exhibited a highly connected solvent network with relatively high *D_f_*, whereas increasing PopZ or Mg^2+^ concentration drove a transition to a core–shell architecture with a water-depleted core, a highly polar shell, and lower *D_f_* values (Supplementary Fig. S3c). This *E_T_* (30)-resolved binning scheme serves as the structural basis by which changes in the area of highly polar pixels (Bin 4), either expansion or contraction, reflect local hydration at the sub-condensate length scales.

**Fig. 2:**
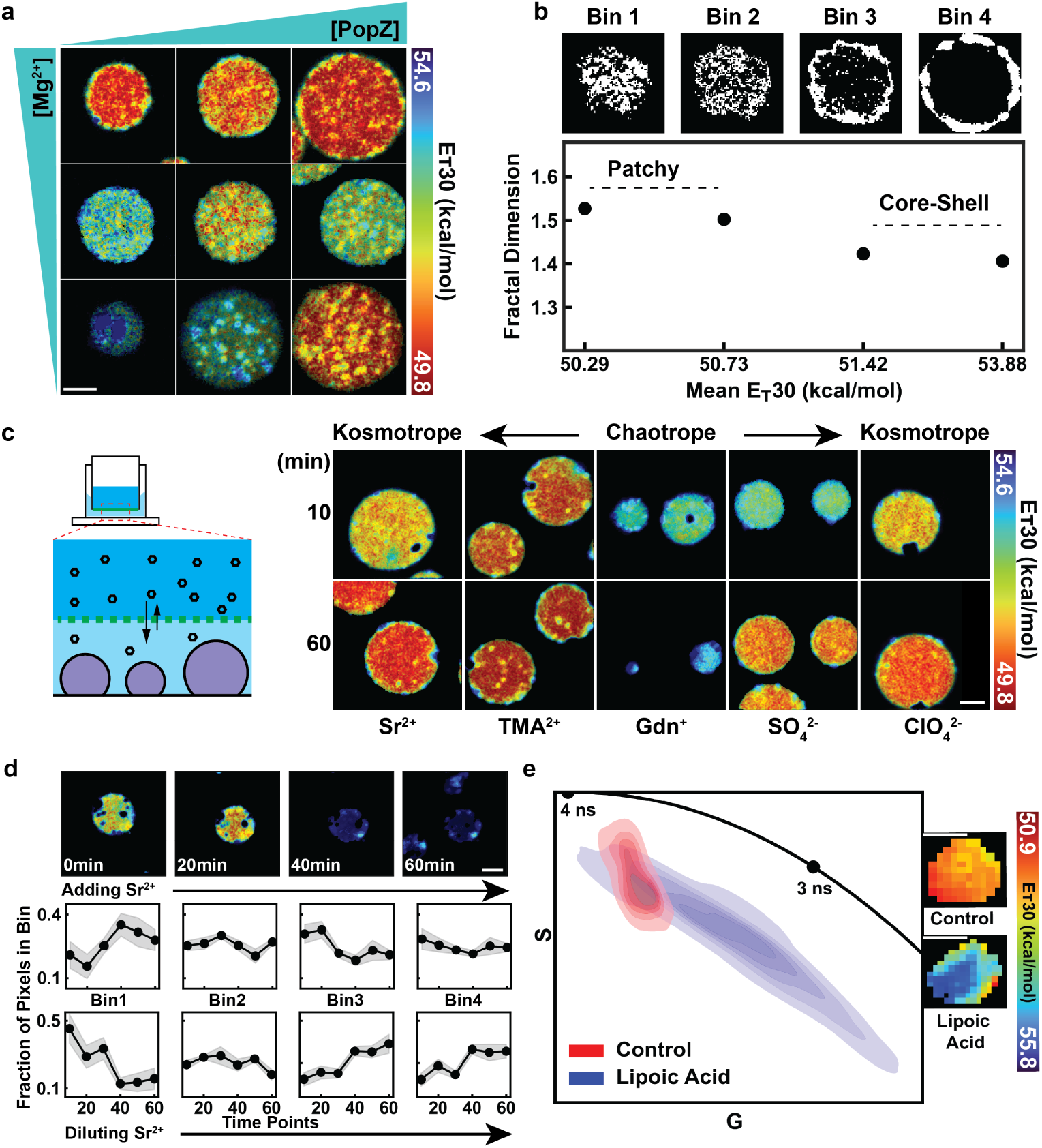
The aqueous architecture of PopZ condensates is a tunable thermodynamic property. **a,** ConSTEL images of PopZ condensates formed at varying concentrations of PopZ and Mg^2+^. Moving further from the phase boundary results in a more hydrophobic core and a distinct polar shell (*n >* 9 condensates per phase-space point from three independent experiments). **b,** Fractal dimension (*D_f_*) of segmented solvent microenvironments. The continuous *E_T_* (30) landscape is thresholded into four polarity bins (Bin 1, most hydrophobic; Bin 4, most aqueous) to quantify spatial connectivity. *D_f_* captures transitions of each microenvironment from an interconnected patchy network to a more isolated architecture. **c,** Left: schematic of the microfluidic dialysis device used for real-time buffer exchange. Right: time-lapse ConSTEL images of PopZ condensates during dialysis with representative kosmotropic (Sr^2+^, TMA^+^, SO_4_^2 –^, ClO_4_ ^−^) and chaotropic (Gdn^+^) ions. Kosmotropes drive water out of the condensates (decreasing *E_T_* (30)), whereas chaotropes drive water in (increasing *E_T_* (30)). **d,** Top: representative time-lapse ConSTEL images during continuous microfluidic dilution of the kosmotrope Sr^2+^. Bottom: fractional area of each polarity bin over a 60 min cycle of Sr^2+^ addition and dilution. The reciprocal reversal of the bin populations shows that architectural tuning tracks Sr^2+^ concentration in a reversible manner, consistent with a fluid thermodynamic equilibrium rather than irreversible kinetic precipitation. Data are mean with shaded s.e.m. (*n* = 9 condensates for Sr^2+^ addition; *n* = 5 for dilution). **e,** Phasor plot and representative ConSTEL maps of Nile Red–labelled PopZ condensates in live *E. coli* following treatment with the dissolution-promoting metabolite lipoic acid or a vehicle control. The shift toward shorter fluorescence lifetimes in phasor space reflects an expansion of the highly polar solvent fraction immediately preceding macroscopic phase collapse. Scale bars, 2 µm for *in vitro* images (c,d) and 500 nm for *in vivo* images (e).

We then examined how chemical stimuli remodel this aqueous architecture. Using a bespoke dialysis platform, we introduced Hofmeister series ions into PopZ condensates in real time during ConSTEL imaging (Fig. 2c). The chaotropic ion guanidinium (Gdn^+^) drove water into the condensate network, manifested as an expansion of Bin 4 at the expense of Bin 1. In contrast, kosmotropic ions such as 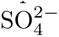, Sr^2+^, 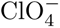, and TMA^+^ compressed the protein-rich scaffold and effectively drove water out. Upon addition of Sr^2+^, the fractional area of Bin 1 rapidly increased while Bins 3 and 4 decreased (Fig. 2c), accompanied by shifts in *D_f_*for each microenvironment (Supplementary Fig. S3d).

A common complication in Hofmeister experiments is irreversible “salting out” or kinetic precipitation of the protein network [39, 40]. To distinguish such arrested states from reversible thermodynamic tuning, we used continuous dialysis to first add and then dilute Sr^2+^ (Fig. 2d). The reciprocal reversal of the polarity-bin fractions upon Sr^2+^ addition and dilution shows that the protein–solvent architecture is responsive and reversible over our observation window, consistent with modulation of a fluid thermodynamic equilibrium rather than solid-state gelation.

Finally, we asked whether this physicochemical control operates in living cells. We perturbed PopZ condensates in live *E. coli* with lipoic acid, a metabolite that has been shown to promote condensate dissolution [14, 27, 41]. Relative to a vehicle control, lipoic acid induced a pronounced shift of the phasor values toward higher *E_T_* (30) values (around 56 kcal mol*^−^*^1^), corresponding to an expansion of a highly polar, Bin 4-equivalent solvent fraction (Fig. 2e). Concurrently, single-condensate morphometry revealed a decrease in circularity and an increase in internal lifetime complexity (Supplementary Fig. S3e). Thus, the internal solvent architecture is a highly plastic material property that can be tuned by ionic composition *in vitro* and by metabolic cues *in vivo*, behaving as a thermodynamic state that links phase behavior to local hydration.

### Spatially Inhomogeneous Network Fluids Compartmentalize Mesoscale Rheology

To determine how the scaffold-solvent architecture shapes mesoscale transport, we combined time-averaged FLIM of Nile Red with single-particle tracking (SPT) of fluorescent nanoparticles (∼142.0 nm diameter, composed of Bodipy-FL C_12_) to simultaneously map the chemical landscape and probe local mesoscale rheology in PopZ condensates (Fig. 3a, S4a-c, Supplementary movie 3). To ensure our measurements reflect true steady-state material properties rather than transient insertion kinetics, SPT was performed after a pre-equilibration. Global phasor analysis confirmed that the fundamental multi-phase heterogeneity of the condensate remained fully intact in this equilibrated state (Fig. S4d).

**Fig. 3:**
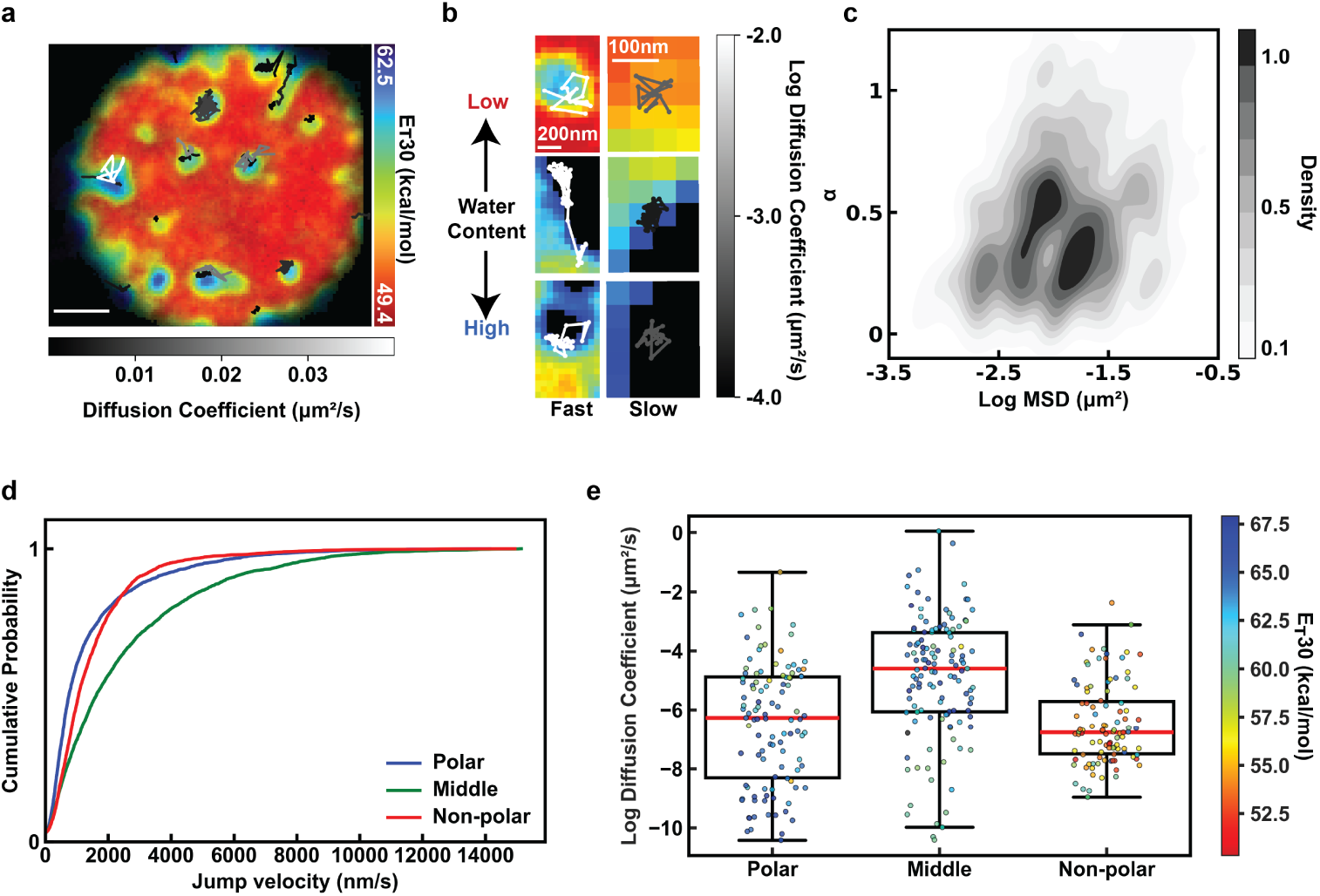
The spatially inhomogeneous network fluid creates distinct rheological compartments. **a,** Representative ConSTEL map of the internal *E_T_* (30) polarity mosaic within a PopZ condensate overlaid with instantaneous nanoparticle trajectories. The vertical colormap denotes local *E_T_* (30) solvent polarity, while the horizontal grayscale colormap designates the apparent diffusion coefficient of the tracers. See Supplementary video 3. **b,** Representative montage of salient nanoparticle trajectories exhibiting fast-to-slow mobility across distinct chemical niches (Polar, Middle, and Non-polar). The grayscale colormap shows the diffusion coefficient, while the vertical double-arrow illustrates relative water content (based on *E_T_* (30) mapping). **c,** Bimodal *D*-versus-*α* (apparent diffusion coefficient versus anomalous diffusion exponent) probability density distribution for the nanoparticle trajectories. The data reveal a spatial decoupling where transport velocity is inversely coupled to spatial confinement (*n* = 389 particles tracked from three independent measurements). **d,** Cumulative jump velocity distributions for nanoparticles segregated by their localized solvent microenvironment (Polar, Middle, and Non-polar). These overarching thermodynamic states were accessed by titrating PopZ and Mg^2+^ concentrations according to the phase boundaries established in Figure 2a. **e,** Box plot showing the distribution of diffusion coefficients across the distinct spatial niches. Each point is color-coded by the average *E_T_* (30) value of the specific pixels traversed by the nanoparticle’s trajectory. Counterintuitively, intermediate polarity domains facilitate the highest mobility via a highly confined ‘cage effect’, whereas thermodynamic extremes (highly Polar and Non-polar domains) act as a dense, high-friction viscoelastic mesh that severely restricts diffusion (*n >* 100 tracks per condition).

For each trajectory, we overlaid its positions onto the lifetime map, assigned an *E_T_* (30) value to every step, and computed the local diffusion coefficient as a function of the solvent polarity encountered along the path. To quantify the resulting rheological landscape, we analyzed the anomalous diffusion exponent (*α*) versus Mean squared displacement (MSD at the third time lag) (Fig. 3c). The data revealed a complex, bimodal distribution of mobilities spanning over two orders of magnitude. Counterintuitively, diffusivity did not scale linearly with maximum hydration. Instead, these populations display a striking topological dependence, where diffusion speed is inversely coupled to spatial confinement.

The first population, localized predominantly within the intermediate-polarity domains, exhibits rapid diffusion (log_10_(*MSD*(*τ* = 3Δ*t*)) ≈ −1.0) but highly sub-diffusive, confined behavior (*α <* 0.5) (Fig. 3d,e, S4e). This paradoxical signature mathematically defines a “cage effect”, whereby nanoparticles rattle at extreme velocities within loose, localized pockets of uncrosslinked solvent, but their long-range movement is ultimately constrained by the surrounding dense protein–solvent. In stark contrast, driving the local environment toward either thermodynamic extremes by increasing macromolecule concentration (Non-polar) or decreasing divalent ion content (Polar), collapsed these cages into a dense, high-friction mesh. The second population, found in the highly Polar (blue) and Non-polar (red) domains, experienced a drop in diffusion (log_10_(*MSD*(*τ* = 3Δ*t*)) ≈ −2.0) accompanied by an increase in *α*, characteristic of slower percolation through a uniform, viscous network.

This physical compartmentalization creates a functional dichotomy: localized structural cages support rapid microscale mixing, whereas the heavily crosslinked viscoelastic mesh restricts global transport. In doing so, PopZ condensates partition their interior into chemically distinct transport niches, suggesting that molecules with different interaction grammars may experience very different effective solubilities and residence times in each niche.

### Local Chemical Environments Direct Intra-Condensate Sorting and Reciprocal Sculpting

We next asked how partitioning into transport niches translates into selective small-molecule localization. If solubility and residence time are set by local *E_T_* (30)-defined niches rather than the droplet as a whole, then guests whose chemical grammar matches a given environment should be enriched there, whereas mismatched guests should be excluded or driven into different regions of the mosaic.

To systematically test how molecular design affects solubility within the *E_T_* (30) mosaic, we introduced a panel of three rationally designed BODIPY derivatives (Fig. 4a, Table S1, Supplementary Note 3) [42, 43]. Although BODIPY emission varies modestly with local environment, these effects are small over the polarity range sampled here and do not alter the qualitative spatial partitioning patterns described below. Probe 1 combines an extended aromatic core (LogP = 5.0) with two terminal hydroxyl (–OH) groups (HBD = 2), providing both a strong drive to partition into the dense phase and robust hydrogen-bonding capacity to engage local aqueous niches. Probe 2 retains the same aromatic core but replaces the hydroxyls with terminal acetate (–OAc) groups (HBD = 0; LogP = 6.5), preserving the partitioning drive while eliminating hydrogen-bond-donor capacity and slightly increasing hydrophobicity. Probe 3 lacks both hydroxyls (HBD = 0) and the extended hydrophobic appendages of Probe 2 (LogP = 3.9), thereby minimizing both dense-phase partitioning and specific hydrogen-bond interactions.

**Fig. 4:**
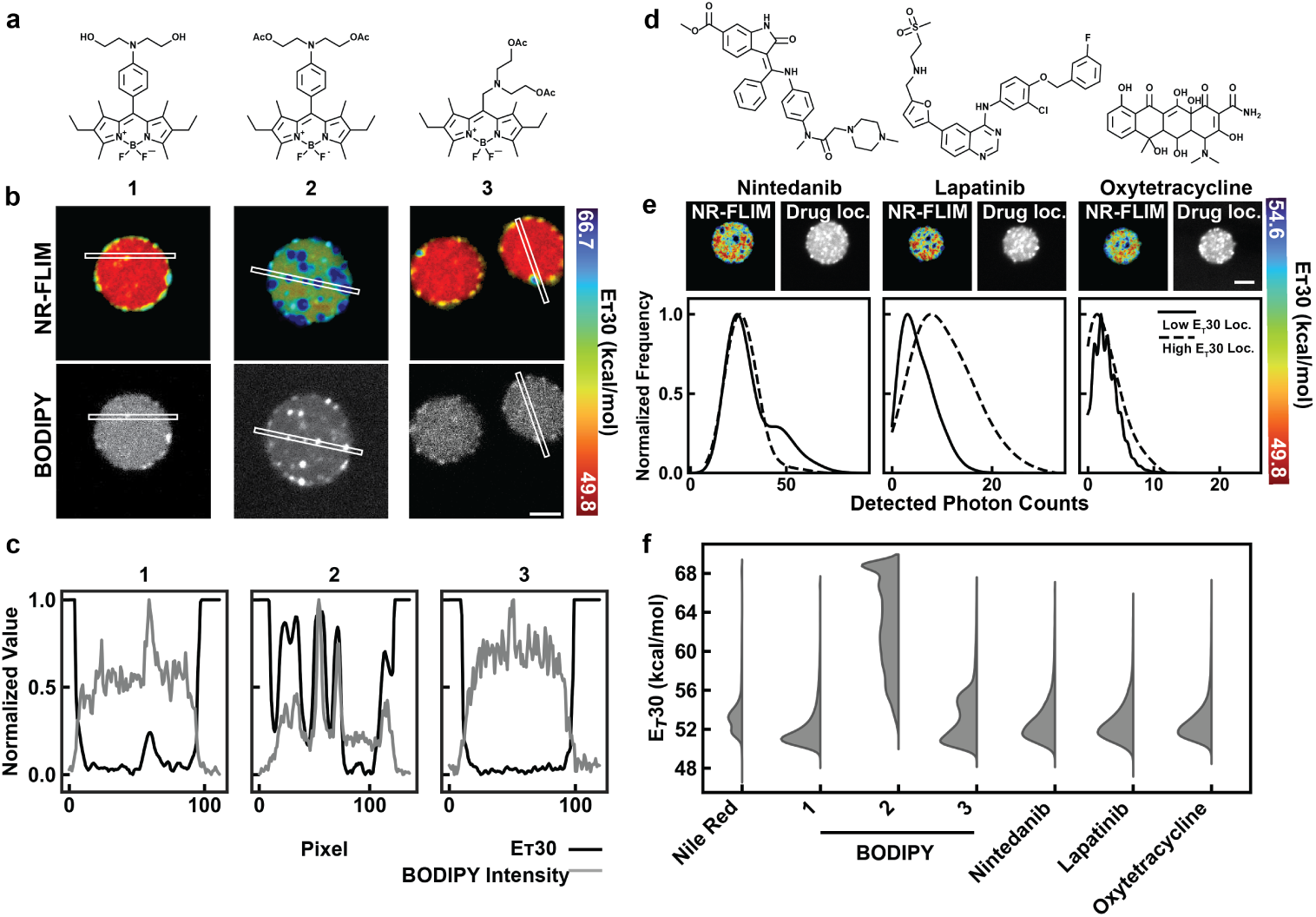
Spatial sorting and reciprocal sculpting in condensate architecture. **a,** Chemical structures of the BODIPY probes. **b,** Representative ConSTEL maps (top) and corresponding probe intensity images (bottom) for each BODIPY molecule. (scale bar is 2 µm) **c,** Line profiles of the *E_T_* (30) polarity (solid) and BODIPY probe intensity (dashed) from the images above. Probes 1 and 3 remain largely diffuse; Probe 2 forms localized aggregates. **d,** Chemical structures of the FDA-approved therapeutics: Nintedanib, Lapatinib, and Oxytetracycline. **e,** Representative ConSTEL maps (left) and corresponding drug localization intensity images (right) (scale bar is 2 µm). Bottom: Corresponding spatial correlation plots showing the normalized frequency of drug photon counts distributed across low and high *E_T_* (30) environments. **f,** Violin plots of global *E_T_* (30) distributions for unperturbed condensates compared to those sequestering probes or therapeutic molecules. Sequestration of the mismatched Probe 2 remodels the environment, demonstrating reciprocal sculpting.

These designs produced distinct patterns of spatial partitioning and local solubility. Probe 1, whose chemical grammar matches both the hydrophobic scaffold and aqueous niches, exhibited the highest partitioning into PopZ condensates (*F_in_/F_out_* ≈ 15) and remained broadly diffuse throughout the network (Fig. 4b,c; Table S1). Probe 3, which retains moderate hydrophobicity but lacks hydrogen-bond donors, showed much weaker overall enrichment (*F_in_/F_out_*≈ 2) and only weak clustering. In contrast, Probe 2 combined strong hydrophobic partitioning (highest LogP) with no hydrogen-bond-donor capacity; it entered the dense phase but rapidly exceeded its local solubility limit, collapsing into severe, highly localized insoluble aggregates despite an intermediate bulk partition ratio (*F_in_/F_out_*≈ 2.4).

We next asked whether therapeutically relevant small molecules navigate this landscape using similar rules. We selected the intrinsically fluorescent FDA-approved drugs Nintedanib, Lapatinib, and Oxytetracycline, which span a wide range of hydrophobicity (LogP from −2.2 to 5.4), hydrogen-bond-donor counts (HBD = 2–7), and polar surface areas (tPSA = 94–202 Å^2^), enabling direct tests of how their chemical grammars map onto the *E_T_* (30) mosaic (Fig. 4d; Table S1). Using detected photon counts as a proxy for local enrichment, spatially correlated pixel analysis revealed that none of the therapeutics distributed uniformly throughout a mean-field hydrophobic phase; instead, each exhibited pronounced localized clustering with niche preferences that mirrored its molecular properties (Fig. 4e; Table S1).

Oxytetracycline, the most polar compound (HBD = 7, tPSA = 202 Å^2^, LogP = −2.2), showed relatively modest overall enrichment but, when it did partition into the dense phase (*F_in_/F_out_* ≈ 4.9), it was concentrated within the highly hydrated, high-*E_T_* (30) regions, consistent with its strong hydrogen-bonding demands. Nintedanib and Lapatinib both feature large hydrophobic architectures dominated by bulky aromatic cores and halogens and accordingly achieve substantial enrichment in PopZ condensates (LogP = 3.2 and 5.4; *F_in_/F_out_* ≈ 2.2 and 4.4, respectively). Despite this similar recruitment by the hydrophobic effect, their local sorting diverges: Nintedanib shows weak clustering with a slight bias toward less polar, scaffold-rich regions (low *E_T_* (30)), whereas Lapatinib forms pronounced clusters within the highly hydrated niches (high *E_T_* (30)). This reveals a spatial decoupling: hydrophobic cores drive thermodynamic recruitment out of bulk solvent, while polar functional motifs and tPSA tune intra-condensate coordinates within the *E_T_* (30) mosaic.

Because the internal *E_T_* (30) mosaic strongly influences a guest’s local saturation limit (*c_sat_*), we next asked whether mismatched molecules can reshape the host architecture. To test this, we quantified the global *E_T_* (30) distributions across titration series for each probe and drug (Fig. 4f and Fig. S5). For Nintedanib, Lapatinib, Oxytetracycline, and the well-matched Probe 1, the condensate solvent landscape remained tightly constrained (∼50–54 kcal/mol) over the full concentration range, indicating non-perturbative integration of these guests. In contrast, Probes 2 and 3 progressively erased the low-*E_T_* (30), less polar patches and drove the interior toward a uniformly high-*E_T_* (30), strongly polar state (∼68 kcal/mol) even at relatively low concentrations. Notably, Probe 1 behaved similarly only at the highest dose: while 5 and 10 µM left the *E_T_* (30) mosaic unchanged, increasing Probe 1 to 50 µM triggered a comparable shift toward this high-*E_T_* (30) state. Thus, matched guests remain soluble and leave the host architecture largely intact across a wide range of concentrations, whereas mismatched or over-saturated guests reciprocally sculpt and ultimately polarize the condensate interior.

Together, these data suggest reciprocal sculpting as a general physical principle in this system: condensate interiors spatially segregate molecules according to local chemical niche, but guests that are fundamentally mismatched to the host network, or that simply exceed local solubility limits, destabilize that architecture. Ultimately, this points to a dual-edged predictive principle for targeted drug design. Because therapeutics undergo localized clustering, rationally matching a drug’s chemical grammar to the context-specific environment of its biological target can be leveraged to amplify its local concentration and efficacy, whereas structural mismatches risk sequestering the drug into non-productive niches or inadvertently disrupting the condensate it aims to occupy.

## Discussion

Recent work has mapped the spatial heterogeneity of condensate scaffolds and characterized solvent states inside and outside dense phases, but the continuous solvent architecture experienced by small molecules within condensates remains unresolved. Here, by applying the solvatochromic dye Nile Red, whose fluorescence lifetime is highly sensitive to hydrogen-bonding interactions with the surrounding solvent, we map this internal architecture and find that the dye is broadly distributed, with apparent intensity “clusters” marking regions of distinct hydrogen-bonding environments rather than discrete scaffold binding sites. Mapping time-averaged lifetimes thus provides direct access to the continuous thermodynamic topology of the internal solvent phase. We reveal that isolated hubs of activity are physically interconnected by a stimulus-responsive aqueous network, supporting the view of biomolecular condensates as microphase-separated, spatially inhomogeneous network fluids.[11, 14, 15, 44, 45] This internal landscape closely mirrors the microphase separation observed in synthetic, non-biological multiphase coacervates,[46–48] suggesting that continuous solvent architectures arise from general Flory–Huggins thermodynamic principles rather than a purely biological anomaly.[49]

This microphase-separated topology provides a structural blueprint for mesoscale rheology and, because its spatial connectivity can be quantified through *E_T_* (30)-resolved fractal dimensions, allows the responsiveness and reversibility of condensate architectures to be measured directly. Correlating local chemical environments with mesoscale transport shows that this topology partitions the condensate into aqueous regions that facilitate rapid local motion and dehydrated regions that dominate long-range mechanical resistance. Crucially, the same solvent-defined niches also modulate chemical reactivity. While current “condenzyme” models attribute catalytic activity primarily to the macromolecular scaffold or macroscopic interfaces,[50, 51] we find that reactions such as ester hydrolysis are neither homogeneously distributed nor fully scaffold-dependent; hydrated solvent niches independently act as nanoscale biochemical reactors.

This internal solvent architecture also exposes a limitation of current models for intra-condensate drug sorting, which rely exclusively on bulk partition coefficients (*C_in_/C_out_*). By averaging over the droplet volume, this metric obscures internal spatial heterogeneity and effectively treats the condensate as a uniformly shared, mean-field hydrophobic phase.[9] Our real-space *E_T_* (30) maps refine this view. By systematically mapping PopZ condensates across their phase diagram, we show that internal solvent chemical environments change markedly with thermodynamic state, so a single condensate can access multiple solvent qualities even at fixed overall composition. Rather than indiscriminate sequestration, thermodynamic recruitment into the macroscopic droplet and subsequent spatial sorting into specific nanoscale coordinates emerge as distinct processes, both governed by local solubility limits within the internal network. In practice, a guest molecule samples different niches according to its polarity, hydrogen-bonding capacity, charge, and hydrophobic surface area, rather than being determined solely by its bulk partition coefficient *C_in_/C_out_*.[4–6] These localized solubility boundaries therefore introduce a physical vulnerability in the condensate architecture, namely reciprocal sculpting. If a therapeutic lacks the required chemical grammar or exceeds its local saturation concentration (*c_sat_*), it thermodynamically displaces the native solvent and disrupts the dense-phase equilibrium. While poorly designed drugs risk inadvertently destabilizing functional biological architectures, this vulnerability also suggests a potential therapeutic strategy. By engineering structural mismatches or overloading the *c_sat_* of specific niches, reciprocal sculpting could be leveraged to thermodynamically dissolve pathological aggregates driving neurodegeneration or oncogenesis.[52, 53] Taken together, our results position condensates as actively tunable solvent architectures rather than passive compartments, and show that resolving their internal chemical microenvironments is essential for predicting, controlling, and deliberately reprogramming molecular function across scales.

## Methods

### PopZ protein purification and condensate sample preparation

PopZ was expressed and purified as previously described [14]. Briefly, PopZ containing an N-terminal His_6_ tag linked via a TEV protease cleavage site was overexpressed in *E. coli* BL21(DE3) and purified under denaturing conditions. Bacterial cells were lysed in a denaturing buffer containing 8 M urea, 500 mM NaCl, 50 mM sodium phosphate (pH 7.0), 10 mM imidazole, and 2 M guanidinium chloride. The His_6_-tagged PopZ was isolated using Ni-NTA resin in a gravity-flow column.

The purified protein was refolded by dialysis and stored at -80 °C in a buffer containing 50 mM sodium phosphate (pH 7.0), 100 mM NaCl, and 10% (v/v) glycerol until further use. The His_6_ tag was cleaved using TEV protease, followed by dialysis into buffer containing 5 mM sodium phosphate (pH 7.0) and 10 mM NaCl prior to condensate preparation. Condensates were formed using the indicated concentrations of PopZ and Mg^2+^ in 5 mM sodium phosphate and 10 mM NaCl buffer (pH 7.0) supplemented with 5 *µ*M Nile Red. Samples were incubated at 30°C for 1 hour before imaging. Prior to adding the condensate samples, the 384-well plate which was incubated with 1% Tween solution for 1 hour at 37 *^◦^*C followed by five washes with water, and subsequently dried under nitrogen gas.[54, 55]

### FLIM setup

Imaging was done on Nikon Eclipse Ti2 microscope equipped with Abberior confocal scanning system. A 100x, NA 1.45 oil-immersion objective and a 1 Airy-unit pinhole was used. The excitation laser and confocal detector outputs were connected to a Time-Correlated Single-Photon Counting (TCSPC) module (Becker & Hickl). For the measurement of Nile Red lifetime, 561 nm excitation pulsed diode laser (40 MHz) at 5.15% power; emission at 575–625 nm was employed. For the measurement of Nile Blue lifetime, 640 nm excitation pulsed diode laser (40 MHz) at 4.95% power; emission at 650-700 nm was employed. All FLIM data were acquired in line-scanning mode (pixel dwell time 10*µ*s) using SPCM Data Acquisition software (v9.89; Becker & Hickl). Unless otherwise noted, fluorescence lifetime images were collected for 30s each.

### Nile Red pure solvent titrations and *E_T_* (30) calculations

A final concentration of 9*µ*M Nile Red was prepared in water, methanol, glycerol, octanol, or a mixture of two solvents immediately before image. Fluorescence lifetime imaging was performed approximately 3*µ*m above the glass surface of a 384-well plate. The acquisition time was adjusted such that the maximum of the photon-count histogram exceeded 100 counts. Fluorescence excitation spectra of Nile Red were recorded for a series of pure solvents with known *E_T_* (30) from [22]. The excitation maximum wavelength for each pure solvent was plotted against its *E_T_* (30) value, and a linear regression was performed to obtain a calibration curve relating excitation maximum to *E_T_* (30) (*R*^2^ = 0.98). For mixed solvents, we first measured the excitation maximum of Nile Red in the mixture solvent and used the calibration line to estimate the effective *E_T_* (30) of the mixture solvent. These effective *E_T_* (30) values for pure and mixed solvents were then plotted against the corresponding fluorescence lifetimes of Nile Red, and a second linear fit was used as a calibration to convert measured lifetimes into effective *E_T_* (30) values in subsequent experiments (*R*^2^ = 0.95).

### Simulations on Nile Red in different environments

All-atom molecular dynamics (MD) simulations were performed to capture the molecular interaction details of Nile Red in distinct chemical environments, including water (*H*_2_O), methanol (MeOH), octanol (OctOH), acetonitrile(ACN), and within a PopZ condensate. For solvent systems, a single Nile Red molecule was randomly placed at the center of a cubic simulation box with side length of 2.5 nm and solvent molecules were randomly inserted to fill up the box. For the condensate system, 14 PopZ protein chains were randomly placed in a 20 nm cubic box and solvated with 249,713 TIP4P water molecules. Then, 24 Nile Red molecules are inserted in the hydrated PopZ phase. Periodic boundary conditions were applied in all three dimensions for all systems. Energy minimization was performed using the steepest descent algorithm, followed by 100 ps equilibration under NVT conditions at 300 K with a 2 fs time step. Subsequent NPT equilibration was carried out for 100 ps using the Parrinello–Rahman barostat to maintain a pressure of 1 bar, releasing all positional restraints. Production simulations were run for 100 ns with a 2 fs time step using GROMACS 2023.3 and the OPLS-AA force field. Energies and coordinates were saved every 10 ps (5,000 steps) for analysis. Hydrogen bonds were identified using a donor–acceptor distance cutoff of 3.2 Å and an angle cutoff of 130*^◦^*. H-bond lifetimes were computed by looping over all frames and tracking continuous bonding periods.

### Time-dependent density functional theory lifetime calculation

TDDFT calculations were performed using ORCA 4.2.1 with the CAM-B3LYP exchange–correlation functional and the 6-31G(d,p) basis set.[56, 57] Solvent effects were incorporated through the conductor-like polarizable continuum model (CPCM). The lowest 10 singlet excited states were included in the TDDFT calculations. Radiative lifetimes were estimated from the S_1_ → S_0_ transition using the calculated oscillator strength and transition energy according to

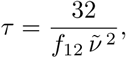

where *f*_12_ is the oscillator strength and 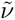 is the transition frequency (in cm*^−^*^1^).

### Ionic modulation through Hofmeister series

A 250 *µ*L sample of condensates containing 5*µ*M PopZ, 100 mM Mg^2+^, and 9 *µ*M Nile Red was added to the imaging chamber. A separate 250 *µ*L solution containing the same components, except for PopZ, was prepared in a 3 kDa cutoff dialysis chamber with the addition of 400 mM of the indicated Hofmeister ion (100mM for Sr^2+^) in sodium phosphate buffer mentioned above. The dialysis chamber was gently lowered into the image cell insert to initiate diffusion of the ion into the condensate system.

For the dilution experiment, a condensate sample containing a final concentration of 50mM Sr^2+^ (250*µ*L total volume) was prepared and incubated for 20 minutes. The sample was then placed in the imaging chamber. A 3kDa cutoff dialysis chamber containing all components except Sr^2+^ and PopZ was carefully introduced into the imaging cell insert to allow diffusion-mediated dilution of Sr^2+^ to from the system. Fluorescence lifetime images of individual condensates were acquired every 10 minutes following the addition of dialysate to the imaging chamber.

### Nanoparticle preparation and correlative tracking with ConSTEL

BODIPY C12 nanoparticles were prepared in 5mM sodium phosphate buffer with 10 mM sodium chloride at pH 7, with a final BODIPY C12 concentration of 50*µ*M. These pre-formed nanoparticles were added to the condensate samples prior to triggering condensate formation with Mg^2+^. Particle tracking within condensates was performed using 561 nm excitation with 5.15% laser power, with emission collected between 575 and 625 nm and a dwell time of 3 *µ*s per pixel, while fluorescence lifetime was recorded concurrently. Time-lapse imaging was acquired at frame intervals ranging from 150 ms to 235 ms. Molecular trajectories were extracted and analyzed using the TrackMate plugin in ImageJ/Fiji. For each trajectory, the mean squared displacement (MSD) was computed as a function of time lag, and the first four MSD points were fitted to obtain the apparent diffusion coefficient (D) and the anomalous diffusion exponent (*α*), derived from; *MSD* = 4*Dt^α^*. For the analysis shown in Fig. 3c, we focused on the short-time MSD at a lag of three frames: for each trajectory we calculated MSD(*τ* =3Δ*t*) from the squared displacements at this lag and paired this value with the corresponding *α* obtained from the per-track fit as mentioned above. The resulting per-trajectory values of MSD(*τ* =3Δ*t*) and *α* were pooled across all movies and used to construct the joint distribution and kernel density estimate of *α* versus MSD(*τ* =3Δ*t*). The resulting tracer trajectories were spatially overlaid onto fluorescence lifetime images, enabling correlation of molecular mobility with local lifetime variations within the same field of view.

### Live *E. coli* Imaging

#### Bacterial Culture and Induction

*E. coli* cells containing the PopZ expression construct were cultured overnight in M9G minimal medium supplemented with kanamycin (30 µg/mL). The following day, cultures were diluted back into fresh M9G medium and grown to an optical density at 600 nm (OD_600_) of 0.35. Protein expression was induced by the addition of isopropyl *β*-D-1-thiogalactopyranoside (IPTG) at a final concentration of 1 mM, and cells were incubated at 37 °C for 24 hours to permit PopZ condensate formation. Following the 24-hour induction, cells were pelleted by centrifugation (15,000 × *g*, 1 min) and resuspended in M9G medium. Electroporation was used to achieve efficient delivery of Nile Red across the bacterial cell envelope and into the cytoplasm, consistent with its established role in enhancing small-molecule 16 and nucleic-acid uptake by transiently permeabilizing membranes.[58] To prepare the cells for electroporation, 500 µL of the suspension was subjected to a two-fold dilution series to standardize cell density, then washed three times with ultrapure water (Milli-Q) to remove salts. A 100 µL aliquot of the diluted cells was transferred to an electroporation cuvette, and Nile Red (18 µM final concentration in dimethyl sulfoxide) was added immediately before a single electrical pulse (1.4 kV, 4 ms) was applied. After electroporation, cells were promptly transferred to Luria–Bertani (LB) medium and incubated at 37 °C for 30 min to allow dye internalization and recovery, then diluted to a final volume of 500 µL with additional LB. Labeled cells were collected by centrifugation (15,000 × *g*, 1 min), washed twice with M9G medium to remove excess Nile Red, resuspended in fresh M9G, and immediately used for imaging. Nile Red-labeled PopZ cells were incubated with lipoic acid (LA) dissolved in dimethyl sulfoxide (DMSO) at a final concentration of 5 µM for 30 minutes at room temperature to allow chemical tuning of the condensate environment. A control experiment was performed in parallel, wherein an equivalent volume of DMSO was added to the cell suspension in place of the lipoic acid stock solution, to account for any effects arising from solvent addition alone. Treated *E.coli* cells were mounted on agar pads prepared from 2% agar dissolved in M9G medium. Samples were imaged at room temperature immediately after mounting. To segment PopZ clusters in E. coli cells (Fig. 1g), we applied intensity thresholding, as the membrane signal intensity was lower than that of the polar PopZ condensates.

### BODIPY derivatives synthesis, characterization, and paritioning assays

See supplementary note 3 for BODIPY derivative syntheses and characterization details, including computed physicochemical properties (Table S1). For the partition assay of the BODIPY-derived probes, condensate samples containing either 10 *µ*M PopZ with 200mM Mg^2+^ or 5 *µ*M with PopZ 50mM Mg^2+^ were used. The fluorescence intensity of BODIPY probes 1,2 and 3, was measured using 488nm excitation at 5.15% laser power, with emission collected between 505-550nm. For Nintedanib, Lapatinib, and Oxytetracycline, condensates containing 5 *µ*M PopZ with 50mM Mg^2+^ were used, and drug fluorescence was measured using excitation at 405nm with 10% laser power and emission collected between at 505-550 nm. In all cases, compounds were added at a final concentration of 5*µ*M immediately before triggering the phase separation, together with 9 *µ*M Nile Red unless otherwise noted. Fluorescence lifetime imaging and intensity scans were performed sequentially.

## Acknowledgements

The authors thank Prof. D. G. Grier (NYU), Prof. P.D. Dahlberg, and Prof. C. Jacobs-Wagner (Stanford University) for helpful discussions. This work was supported in part through the NYU IT High Performance Computing resources, services, and staff expertise. This study was funded by Simons Center for Computational Physical Chemistry at NYU (SF Grant No. 839534) to NH, a U.S. DOE, Office of Science, BES (DE-SC0026328) to NH, and a National Institutes of Health award (1R35GM157103) to SS. High-resolution mass spectra were recorded in the Department of Structural Studies of the N. D. Zelinsky Institute of Organic Chemistry, Moscow. The xSight instrument used for holography was acquired as shared instrumentation with support from the MRSEC program of the NSF under award DMR-1420073.

## Declarations

M.S. and S.S. are inventors on pending patent applications related to the ConSTEL methodology described in this work. The other authors declare no competing interests.

All authors consent to publication.

Data and Materials are available on request.

All code used in this study is available at github.com/saurabhLabNYU/ConSTEL.

Author contributions. S.S. and M.S. conceived the study. S.S. supervised experiments, analyses and implementation of the FLIM setup. M.S., S.B., R.Z., and J.v.H. performed experiments. Y.V. and U.D. synthesized compounds. M.C., N.H., and J.H. performed computations. S.S. and M.S. wrote the manuscript with input from all authors.

## Extended Data

**Fig. S1:**
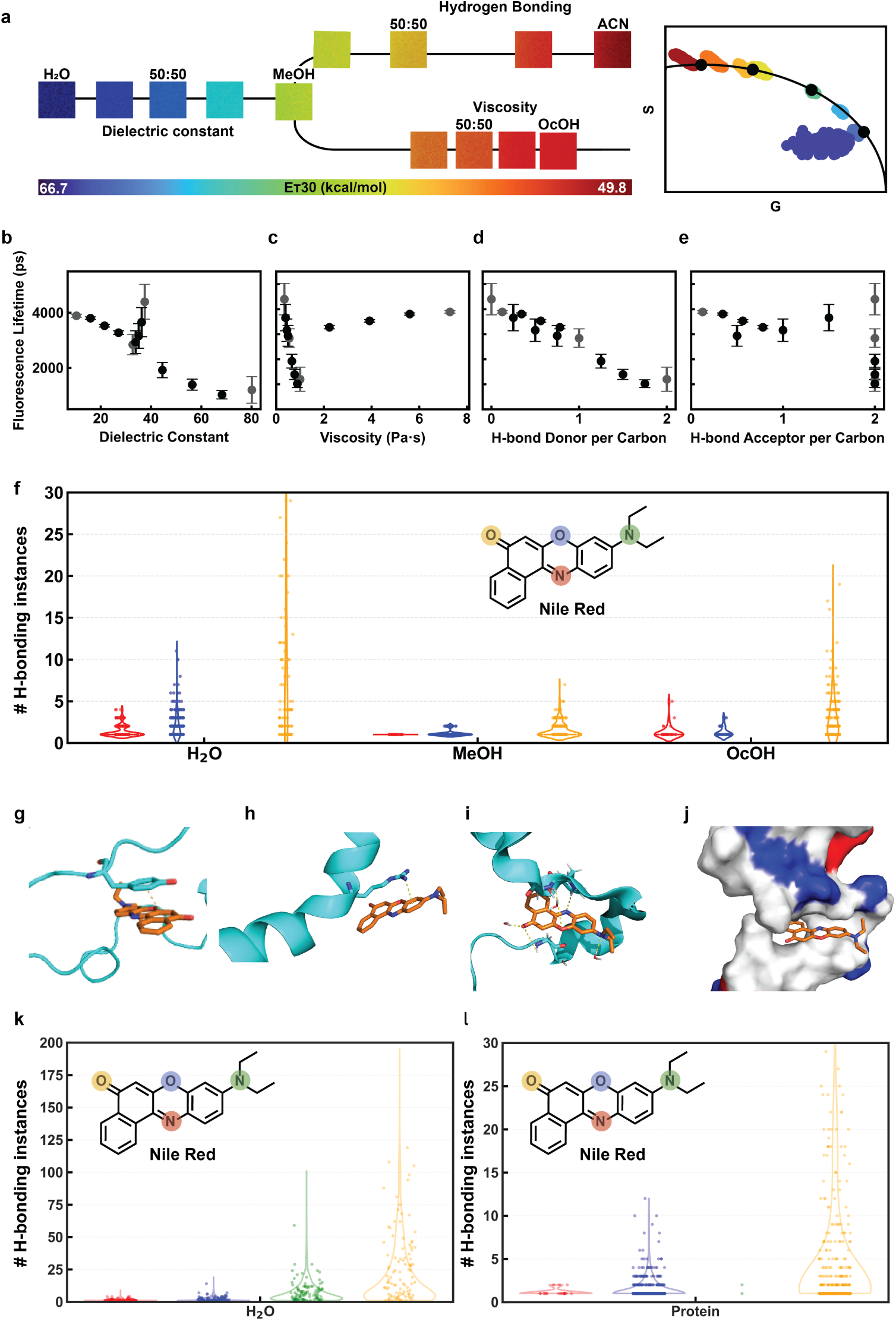
Nile Red fluorescence lifetime is a robust sensor of the local hydrogen-bonding. **a,** Fluorescence emission lifetime images of Nile Red in binary solvent mixtures of methanol/water, acetonitrile/methanol, and octanol/methanol allow for a systematic exploration of solvent parameters—dielectric constant, hydrogen bonding, and viscosity, respectively—on the lifetime response of Nile Red. The color scale represents the empirical polarity parameter *E_T_* (30). Corresponding phasor plot analysis (right) illustrates the trajectory of the Nile Red population across diverse chemical environments (representative dataset of phasor plot from two independent experiments, *n >* 2000 pixels). **b–e,** Plots showing the relationship between Nile Red fluorescence lifetime and bulk solvent properties: **b**, dielectric constant **c**, viscosity **d**, hydrogen-bond donor capacity per carbon **e**, and hydrogen-bond acceptor capacity per carbon. Lifetime correlates most strongly with the hydrogen-bond donor capacity of the solvent. Data are presented as mean ± s.d. (representative dataset from two independent experiments, each with *n >* 10, 000 pixels). **f,** Distribution of solvent–Nile Red hydrogen-bonding events from MD simulations for water, methanol, and octanol. The colored heteroatoms on the Nile Red structure correspond to the same color code in the violin plots, illustrating which sites on the dye most frequently participate in hydrogen bonding in each solvent. **g–j,** Representative snapshots of Nile Red within a simulated PopZ condensate, depicting *π*-*π*, cation-*π*, water-mediated H-bonds, and hydrophobic molecular interactions. Also see SI video S2. **k,** Violin plots of *water-mediated hydrogen bond interactions* with different color-coded heteroatoms in Nile Red based on MD simulations in a PopZ dense phase. **l,** Violin plots of *protein-mediated hydrogen bond interactions* with different color-coded heteroatoms in Nile Red based on MD simulations in a PopZ dense phase. Note the difference in y-axis ranges in **k** and **l**.

**Fig. S2:**
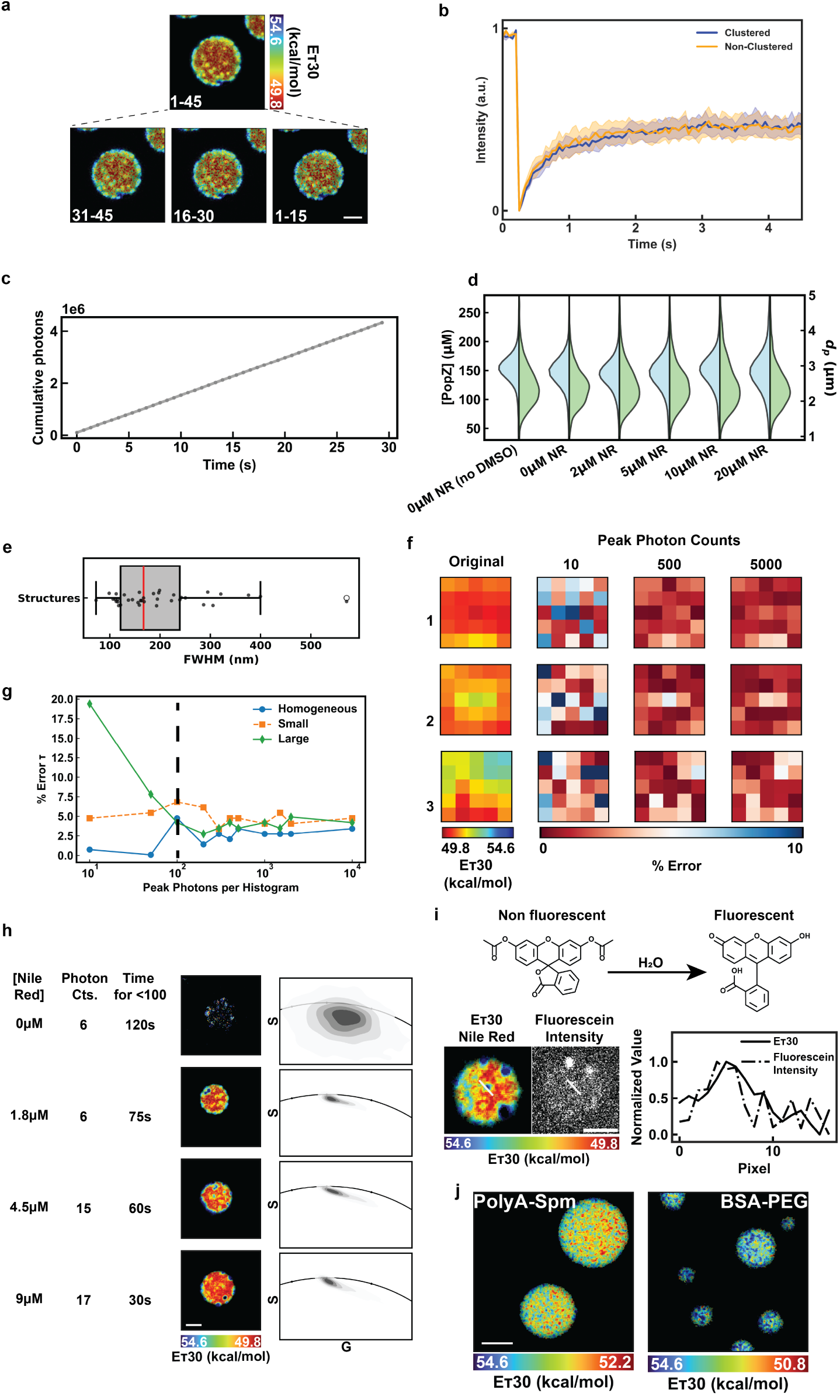
Photophysical validation, continuous stability, and functional mapping of the ConSTEL approach. **a,** Verification of structural persistence, demonstrating that the spatial *E_T_* (30) profile remains completely stable over a 45-slice continuous acquisition window, (scale bar = 2 µm). **b,** Fluorescence recovery after photobleaching (FRAP) dynamics of Nile Red within the highly intense structural clusters versus non-clustering regions. The comparable recovery dynamics confirm similar molecular mobility, demonstrating these polar clusters are dynamic liquids rather than dye aggregation artifacts. **c,** Validation of Nile Red as a passive, non-perturbative probe. The linear increase in cumulative photon counts over the 30 s measurement window and, **d,** the absence of detectable alterations in condensate size or internal PopZ concentration up to 20 µM Nile Red measured by label-free, attachment-free holography ensure the mapped architectures reflect equilibrium states. **e,** Size distribution (FWHM in nm) of the discrete spatial structures visualized via the polarity map, predominantly falling within the 100–300 nm size range. **f,** PSF-mixing Monte Carlo FLIM simulation with Poisson noise and subsequent grid-based maximum likelihood estimation of lifetimes to determine error. **g,** Error quantification confirms that obtaining *>*100 detected photons per pixel at the photon arrival time histogram peak yields a minimal lifetime estimate error of ∼5%. **h,** Optimization of imaging conditions to ensure *>*100 photon counts at the histogram peak within a 30 s acquisition period, establishing 9 µM Nile Red as the optimal concentration. (scale bar = 2 µm) **i,** Chemical reaction schematic of the water-dependent hydrolysis of non-fluorescent fluorescein diacetate (FDA) to fluorescent fluorescein. A representative ConSTEL map of the solvent architecture (left) and the corresponding fluorescein intensity image (right) demonstrate that functional reactivity is spatially co-localized within the highly polar microenvironments. Line profiles along the indicated white lines show the *E_T_* (30) values (solid) and fluorescein intensity (dashed). (scale bar = 2 µm) **j,** Validation of the ConSTEL approach in alternative condensate systems, including PolyA–Spermine complex coacervates and BSA–PEG folded-protein condensates, demonstrating the broader applicability of mapping nanoscale polarity differences in spatially inhomogeneous network fluids. (scale bar = 2 µm)

**Fig. S3:**
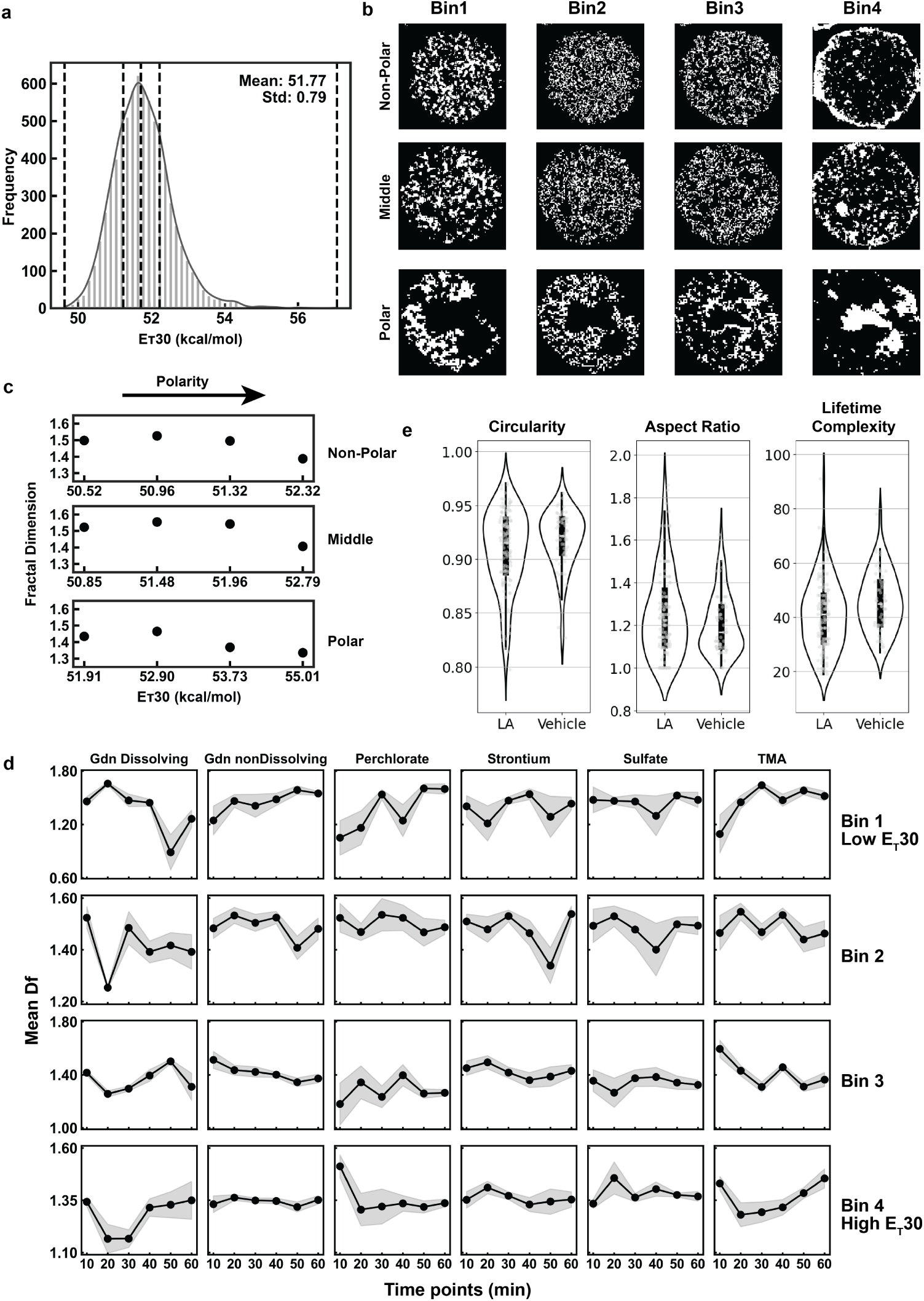
Topological quantification and morphometric analysis of condensate architectures. **a,** Global frequency distribution of *E_T_* (30) values across a representative condensate population. The dashed lines explicitly define the quantitative *E_T_* (30) thresholds used to segment the optical maps into the four discrete thermodynamic bins. **b,** Binary segmentation maps of the spatial polarity mosaic into discrete thermodynamic bins (Bin 1 to Bin 4) for condensates exhibiting non-polar, middle, and polar average architectures. **c,** Calculated fractal dimensions (*D_f_*) of the internal network as a function of mean *E_T_* (30) polarity. Condensates near the phase boundary (Polar and middle) exhibit a highly connected ‘patchy’ state, whereas those deeper in the two-phase region (Non-Polar) transition into a disconnected ‘core-shell’ state. **d,** Changes in fractal dimension (*D_f_*) over time for each polarity bin during microfluidic dialysis with kosmotropic and chaotropic ions. Data represent more than seven condensates measured across two independent experiments (shaded regions represent s.e.m.). **e,** Single-condensate morphometric analysis (circularity, aspect ratio, and lifetime complexity) of PopZ condensates observed in live *E. coli* cells treated with lipoic acid (LA) versus vehicle control, demonstrating structural deformation alongside thermodynamic tuning.

**Fig. S4:**
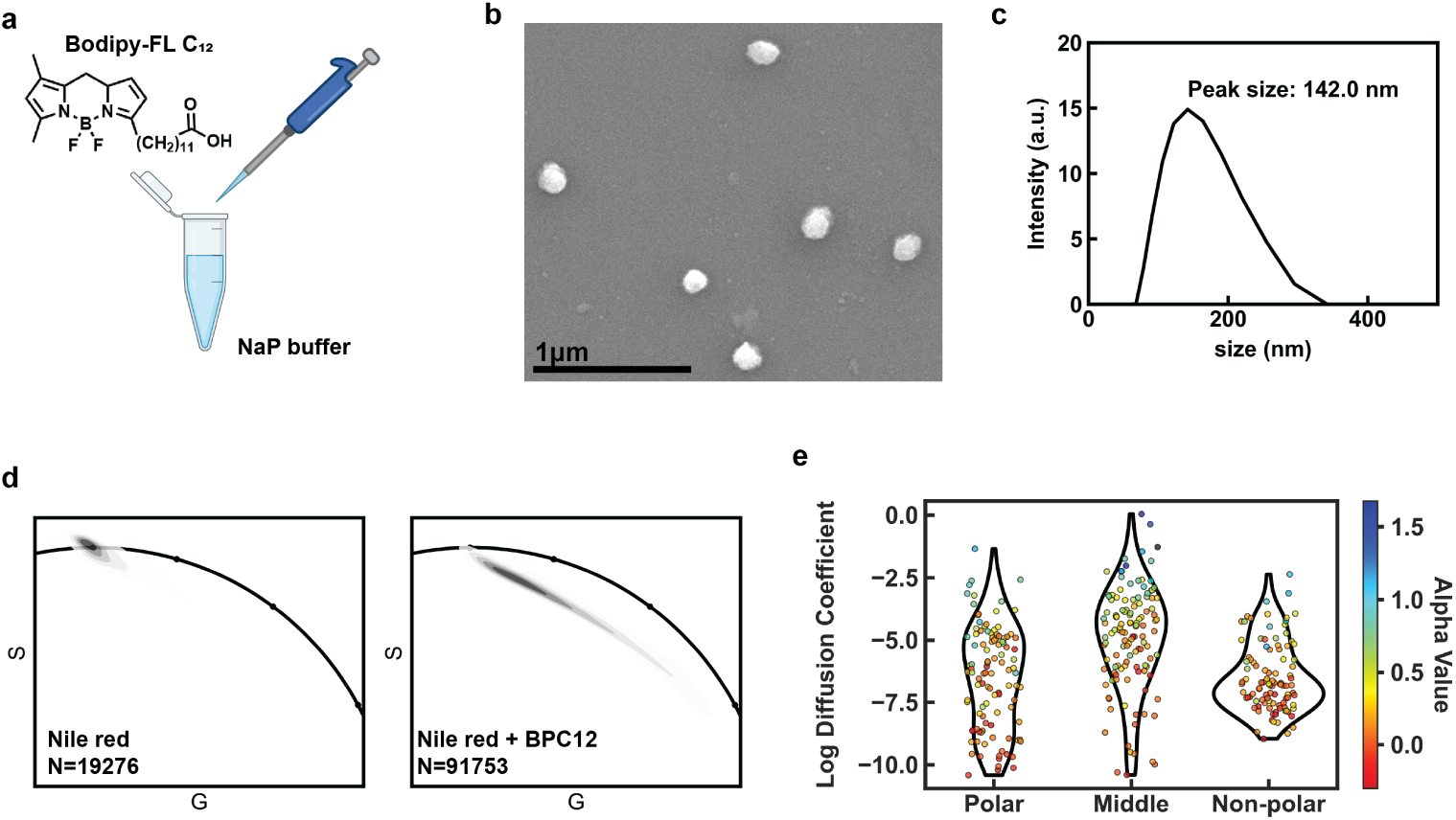
Nanoparticle characterization and mesoscale rheology within PopZ condensates. **a,** Schematic of BODIPY-FL C_12_ nanoparticle preparation. NaP is Sodium phosphate buffer. **b,** Representative scanning electron microscopy (SEM) image of the BODIPY-FL C_12_ nanoparticles. **c,** Representative size distribution of the nanoparticles, confirming a peak diameter of 142.0 nm from two independent measurements. **d,** Global phasor plot analysis showing the time-domain fluorescence lifetime populations for condensates labeled with Nile Red alone (left) and following a 1-hour equilibration with the nanoparticles (right). Insertion of mesoscale cargo leads to higher polarity in PopZ condensates. The number of pixels analyzed (N) is indicated in the respective graphs. **e,** Violin plots showing the distribution of the log value of the apparent diffusion coefficient (*µ*m^2^*/*s*^α^*) of nanoparticles across distinct local chemical microenvironments (Polar, Middle, and Non-polar). The individual data points are color-coded by their corresponding anomalous diffusion exponent (*α*), illustrating the inverse coupling between transport velocity and spatial confinement. (*n >* 100 particles tracked for each condition).

**Fig. S5:**
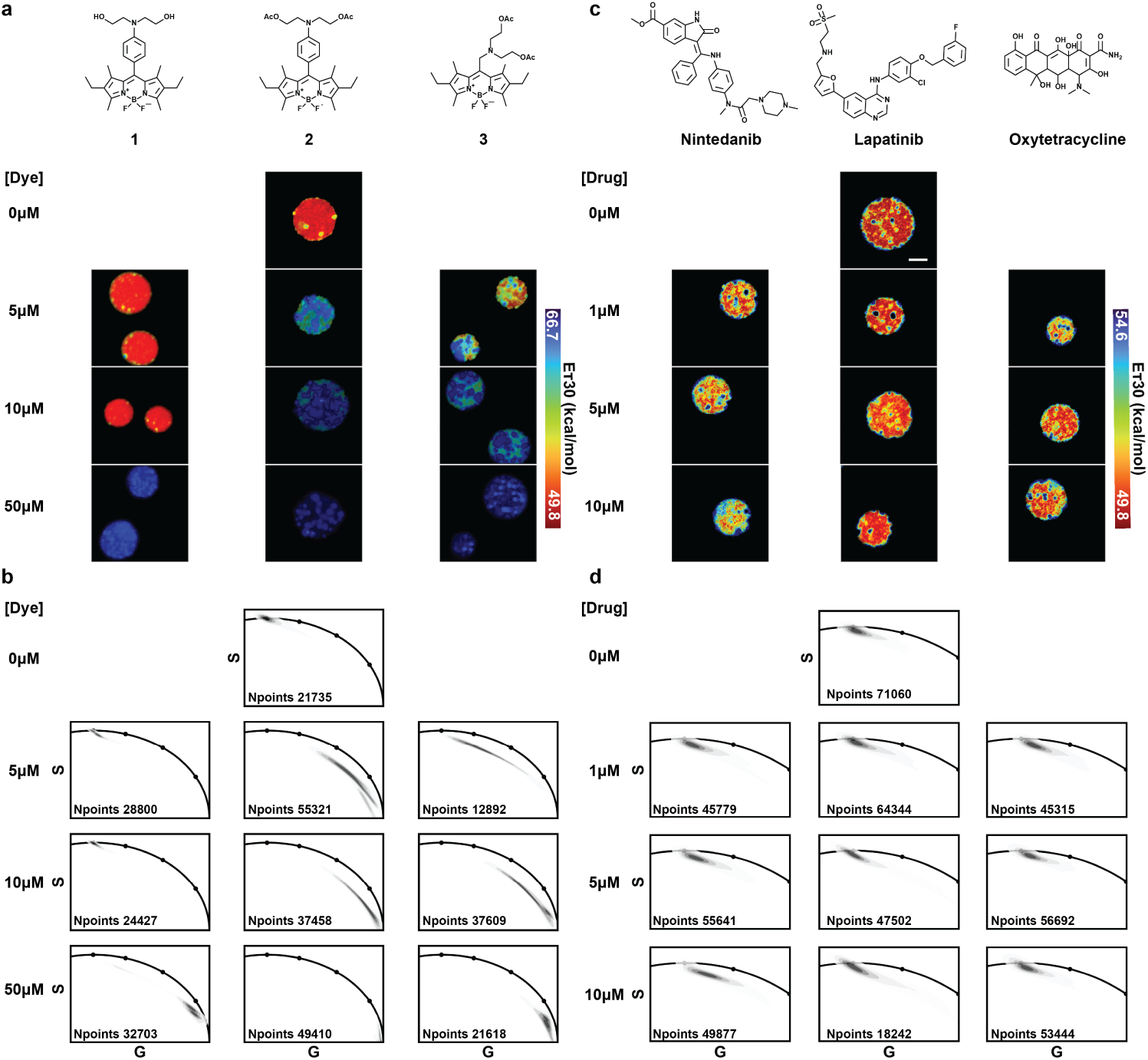
Thermodynamic mapping of the solvent architecture in the presence of guest molecules. **a**, Representative ConSTEL *E_T_* (30) maps of PopZ condensates sequestering BODIPY probes 1–3 at the indicated concentrations, with corresponding chemical structures shown above. **b**, Phasor plots for the conditions in **a**, reporting the distribution of Nile Red lifetimes; the total number of pixels analyzed (N) is indicated in each panel. **c**, Representative ConSTEL *E_T_* (30) maps of condensates sequestering the intrinsically fluorescent therapeutics Nintedanib, Lapatinib, and Oxytetracycline at the indicated concentrations, with chemical structures shown above. **d**, Phasor plots for the conditions in **c**, showing the associated Nile Red lifetime distributions and pixel counts. Scale bar, 2 µm.

## Supplementary Note 1 - Environmental sensing by Nile Red

Dipole–*π* interactions, which generally occur between polar solvents and Nile Red (NR), represent a weak electrostatic attraction between the solvent’s permanent dipole and the dye’s electron-rich aromatic system. TDDFT calculations performed in ORCA (CAM-B3LYP functional and 6-31G(d,p) basis-set) provided the following dipole–*π* interaction energies: acetonitrile = *−*0.556 kcal/mol, methanol = *−*1.839 kcal/mol, water = *−*1.838 kcal/mol, and octanol = *−*2.155 kcal/mol. The negative values indicate that these interactions help stabilize the *π*-electron cloud on the aromatic rings, which will suppress nonradiative decay pathways and lead to a slightly longer fluorescence lifetime. Although acetonitrile exhibits the smallest magnitude of dipole–*π* stabilization, this interaction becomes sufficient to prolong the fluorescence lifetime when hydrogen-bond quenching is absent.

In protic solvents such as methanol or water, however, hydrogen bonding becomes the dominant interaction, polarizing the carbonyl group and drawing electron density toward the carbonyl oxygen. Hydrogen-bond interaction energies are calculated via TDDFT (ORCA: CAM-B3LYP functional and 6-31G(d,p) basis-set)), are: methanol = *−*6.479 kcal/mol, octanol = *−*6.902 kcal/mol, and water = *−*7.2497 kcal/mol, demonstrating the stronger stabilization provided by protic solvents compared to dipole–*π* interactions. These hydrogen bonds act as efficient quenchers by enhancing the rate of nonradiative decay through vibrational coupling pathways that connect the excited electronic state to solvent motions. As a strong hydrogen-bond acceptor, Nile Red forms transient H-bonds that stabilize its carbonyl lone pairs and lower the excited-state energy, effectively reducing the *S*_1_–*S*_0_ gap and facilitating internal conversion. Consequently, fluorescence lifetimes decrease progressively from octanol to methanol to water, directly reflecting the increasing strength and frequency of hydrogen bonding. The exceptionally long lifetime observed in ACN can thus be rationalized by its lack of hydrogen-bond donors, despite its high polarity. This result highlights that hydrogen bonding, rather than polarity alone, is the principal factor governing Nile Red’s excited-state dynamics. These distinct interaction modes are illustrated by TDDFT-calculated electrostatic potential maps of Nile Red in representative pure solvents (Supplementary Fig. S6), where solvent molecules either engage the aromatic core via dipole–*π* contacts or form direct hydrogen bonds to the carbonyl oxygen.

Simulations further reveal that solvent size and steric accessibility are critical to Nile Red fluorescence lifetime. This is particularly relevant when considering the long-established Twisted Intramolecular Charge-Transfer (TICT) model, which has historically been used to explain the photophysics of Nile Red. The TICT theory suggests the twisting of the diethylamino group, a motion that is highly dependent on the amount of empty or “free volume” in the surrounding solvent matrix,

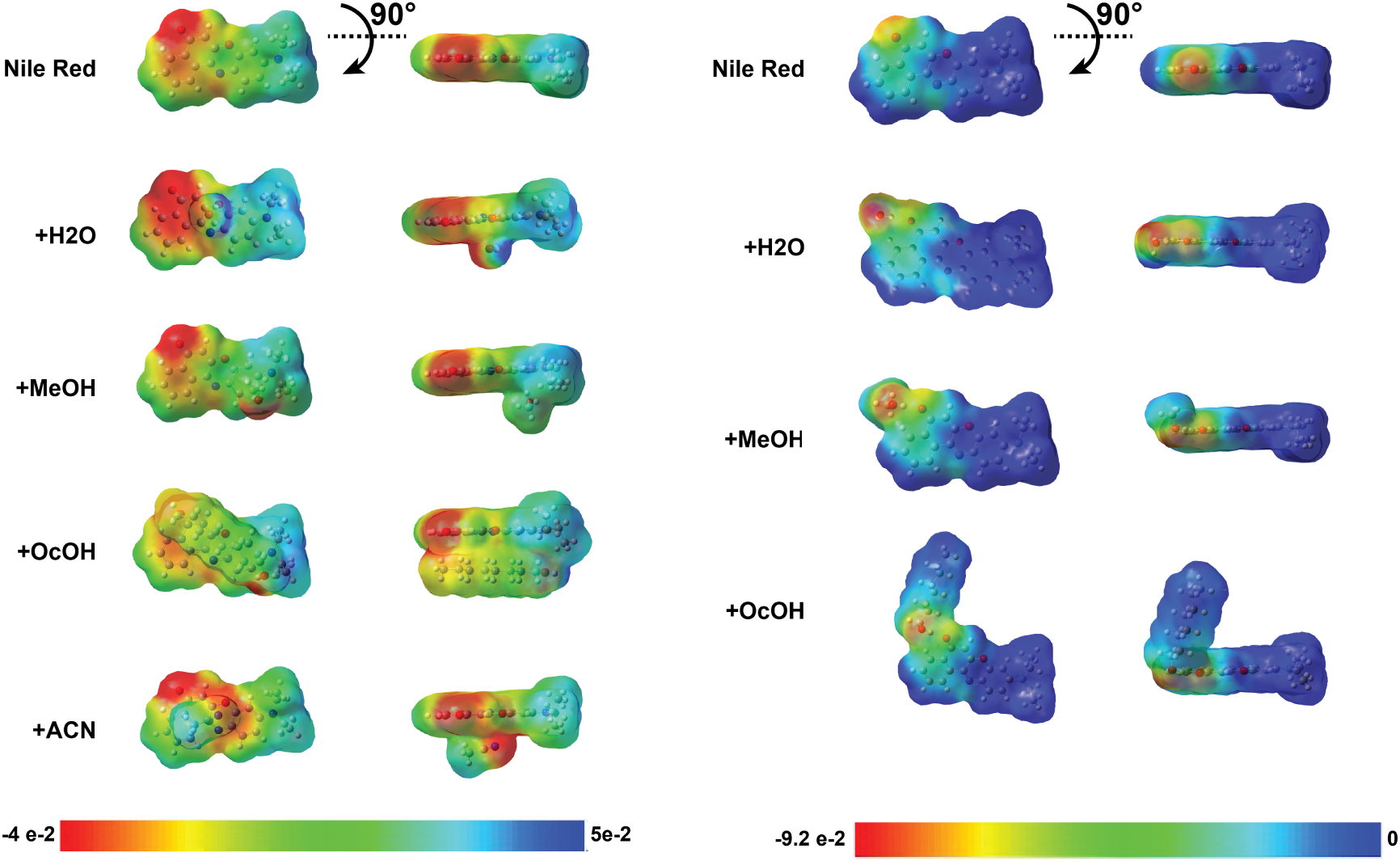

### Electrostatic potential maps of Nile Red in pure solvents

Electrostatic potential (ESP) surfaces of Nile Red caclulated by TDDFT in ORCA for acetonitrile, methanol, water, and octanol, illustrating how different pure solvents redistribute electron density around the carbonyl and amino groups and thereby modulate dipole–*π* and hydrogen-bonding interactions that control the excited-state lifetime. (Left) ESPs of Nile Red and pure solvent structures where the solvent interacts with the aromatic core, vs (Right) the Carbonyl oxygen to be a primary nonradiative decay pathway for Nile Red. When applying this free-volume theory to methanol and octanol, a clear contradiction emerges. Octanol, with its long hydrocarbon tails, has significantly more free volume than the smaller, more efficiently packed methanol molecules. According to a strict interpretation of the TICT model, the greater free volume in octanol should allow for easier twisting of the diethylamino group on Nile Red. Once TICT is enhanced, a nonradiative decay pathway is opened, leading to a shorter fluorescence lifetime in octanol. However, our experimental results show the exact opposite. This finding compellingly disproves the TICT model as the dominant factor for Nile Red in these solvents.

Instead, our results indicate that the fluorescence lifetime of Nile Red is dominated by the efficiency of hydrogen-bond-mediated quenching, which is sterically controlled by the size and dynamics of the solvent. Water, being the smallest and the most flexible, rapidly reorients to form multiple transient hydrogen bonds with NR, strongly enhancing nonradiative decay of Nile Red and leading to the shortest lifetimes. Methanol, though also an effective donor, is larger and less flexible, leading to slightly reduced quenching efficiency relative to water. In contrast, bulky solvents such as octanol are sterically hindered from forming geometrically optimal H-bonds and reorient far more slowly. This suppression of efficient H-bonding interactions diminishes nonradiative decay pathways, yielding longer lifetimes that approach those observed in the aprotic ACN environment.

Having established the primacy of hydrogen bonding in these simple solvent systems, we next sought to determine if this principle holds true within the complex and heterogeneous environment of a PopZ biomolecular condensate. Inside the condensate, nile red is exposed to a variety of potential non-covalent interactions with the protein scaffold, each capable of influencing its local environment and photophysical response. These include *π*–*π* stacking, cation–*π* interactions, and hydrophobic packing.

The aromatic system of nile red and the aromatic side chains of residues like phenylalanine and tyrosine can form *π*–*π* stacking. The strength of the interaction is governed by a balance of electrostatic and dispersion forces. By physically constraining the dye, this stacking can reduce nonradiative decay pathways associated with molecular motion, leading to a longer fluorescence lifetime.

A second powerful non-covalent force is the cation–*π* interaction, an electrostatic attraction between a positive charge from a protonated residue (e.g., lysine, arginine) and the negative potential face of the aromatic system of Nile Red. This interaction can directly alter the electronic structure of the *π* system of Nile Red, thereby influencing its fluorescent properties. By electrostatically stabilizing the *π*-electron cloud, this interaction makes the excited state less susceptible to deactivation through vibrational pathways, leading to a reduction in the rate of nonradiative decay and a longer fluorescence lifetime.

Hydrophobic packing, driven by the thermodynamic drive to minimize contacts between nonpolar groups and water, can sequester Nile Red within less polar, sterically confined pockets of the condensate. Such shielding from water molecules, highly efficient fluorescence quenchers, is therefore expected to increase the fluorescence lifetime of Nile Red locally.

While all of these interactions are possible, our simulations indicate that hydrogen bonding is the most prevalent and dynamically persistent interaction type, occurring continuously and exerting the greatest influence on the system. In the PopZ box solvated with water, hydrogen bonding is overwhelmingly dominated by water, with 66,024 frames (96.5%) arising from H_2_O and only 2,360 frames (3.5%) from protein interactions. For water, the dominant acceptor sites on Nile Red are the carbonyl oxygen with 24,573 frames (37.22%), followed by the diethylamino nitrogen with 23,070 frames (34.94%), the oxazine ring oxygen with 11,344 frames (17.18%), and finally the oxazine ring nitrogen with 7,037 frames (10.66%). This indicates that water molecules most efficiently hydrogen bond with the carbonyl oxygen and the tertiary amine substituent, consistent with their high accessibility and polarity.

In contrast, protein interactions are both rarer and more selective. The carbonyl oxygen accounts for the majority of protein hydrogen bonds with 1,713 frames (72.58%), followed by the oxazine ring oxygen with 622 frames (26.36%), while the oxazine ring nitrogen contributes minimally (22 frames, 0.93%) and the diethylamino nitrogen is almost entirely inaccessible (3 frames, 0.13%). This pattern highlights the strong steric hindrance around the diethylamino substituent, which makes it essentially unreachable for protein donors, whereas the exposed carbonyl oxygen remains the primary locus of protein–dye hydrogen bonding.

This finding demonstrates that even within a protein condensate where *π*–*π*, cation–*π*, and hydrophobic interactions are possible, nile red is not primarily probing the protein itself but is instead reporting on the local aqueous environment. The principle of H-bond-mediated quenching is therefore the critical parameter to interpret Nile Red’s fluorescence lifetime.

## Supplementary Note 2 – Non-perturbative validation and optical controls of ConSTEL

To ensure that the nanoscale polarity mosaics mapped by ConSTEL represent native thermodynamic states rather than probe-induced or optical artifacts, we systematically validated the chemical, dynamic, and optical passivity of the assay.

### Dynamic stability and exchange

Time-averaged fluorescence lifetime imaging microscopy requires continuous laser scanning, raising the potential for photobleaching, local heating, or the artificial optical induction of spatial features. We evaluated probe stability by photon collection over a 45-slices continuous acquisition window (**Supplementary Fig. S2a**). The linear accumulation of photons over time confirms the absence of rapid photobleaching and indicates that the probe is not irreversibly sequestered into static aggregates, but rather exchanges continuously with the bulk solvent in a dynamic equilibrium (**Supplementary Fig. S2b,c**).

### Thermodynamic and chemical passivity

Introducing a solvatochromic guest into a delicate coacervate network carries the theoretical risk of altering thermodynamic tie lines, driving artificial polymer crosslinking, or shifting the binodal phase boundary. To evaluate this, we performed high-throughput quantitative holographic analysis across a broad concentration gradient of the probe. Titrating Nile Red up to 20 µM—a concentration well above our standard experimental dosage of 5–9 µM—altered neither the dense-phase protein concentration ([PopZ]) nor the macroscopic droplet size distribution (*d_p_*) (**Supplementary Fig. S2d**). These data indicate that the probe does not detectably shift the thermodynamic phase equilibrium and acts as a chemically passive reporter.

### Optical sampling and spatial oversampling

The discrete solvent microenvironments exhibit spatial dimensions in the 100–250 nm range (**Supplementary Fig. S2e**), placing them at or near the Abbe diffraction limit of the imaging system. To avoid under-sampling these features, we adopted a spatial sampling grid substantially finer than the diffraction-limited point-spread function by using 50 nm pixels at a 10 µs dwell time. Spatial downsampling controls demonstrated that coarser sampling compresses local time-domain decays into an apparent single-component response, effectively enforcing a bulk ensemble average that masks the multicomponent complexity of the network fluid. In contrast, the oversampled condition preserves the spatially varying lifetime signatures associated with distinct solvent microenvironments.

### Statistical resolution and Monte Carlo error mapping

To determine the photon statistics required to accurately extract lifetimes from these localized volumes, we performed Monte Carlo FLIM simulations. By introducing Poisson noise to simulated decays spanning the full range of observed lifetimes, we mapped the estimation error as a function of peak photon counts (**Supplementary Fig. S2f**). This analysis established a practical threshold: *>*100 detected photons at the histogram peak are required to maintain lifetime estimation error at *∼*5% (**Supplementary Fig. S2g**). Guided by this threshold, experimental acquisition parameters were optimized (yielding 9 µM Nile Red as the ideal concentration) to ensure sufficient photon counts without exceeding the thermodynamic boundaries established in the holography controls (**Supplementary Fig. S2h**). Subsequent systematic reductions in scan iterations and laser power degraded the apparent spatial complexity and increased the inferred lifetime variance, consistent with the emergence of low-photon shot noise rather than a change in the underlying material state.

## Supplementary Note 3 - Synthesis and characterization of small molecules

**Table S1.**
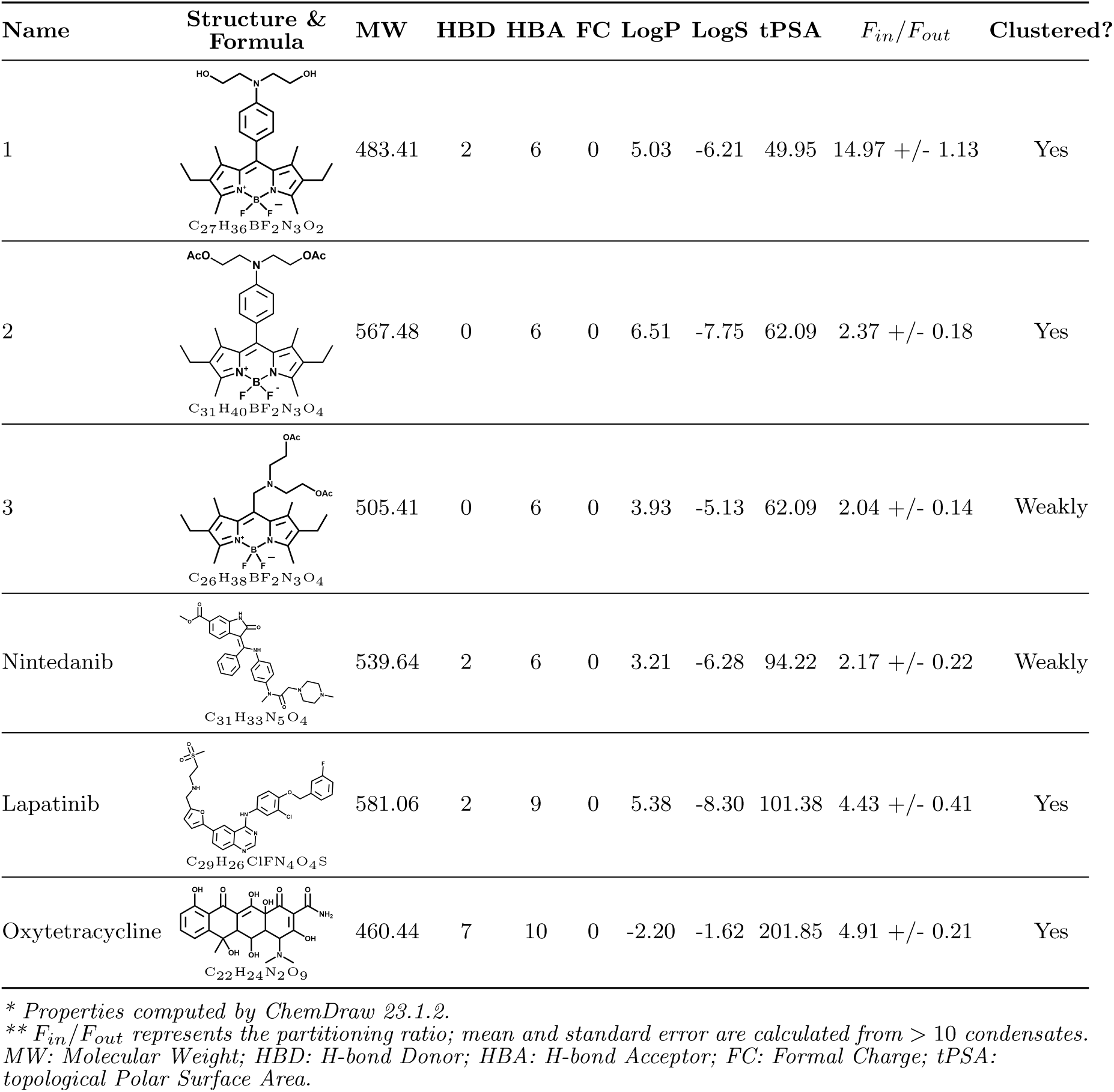
Molecular properties of investigated compounds and reference drugs.

| Name | Structure & Formula | MW | HBD | HBA | FC | LogP | LogS | tPSA | $F_{in}/F_{out}$ | Clustered? |
| --- | --- | --- | --- | --- | --- | --- | --- | --- | --- | --- |
| 1 | C27H36BF2N3O2 | 483.41 | 2 | 6 | 0 | 5.03 | -6.21 | 49.95 | 14.97 +/- 1.13 | Yes |
| 2 | C31H40BF2N3O4 | 567.48 | 0 | 6 | 0 | 6.51 | -7.75 | 62.09 | 2.37 +/- 0.18 | Yes |
| 3 | C26H38BF2N3O4 | 505.41 | 0 | 6 | 0 | 3.93 | -5.13 | 62.09 | 2.04 +/- 0.14 | Weakly |
| Nintedanib | C31H33N5O4 | 539.64 | 2 | 6 | 0 | 3.21 | -6.28 | 94.22 | 2.17 +/- 0.22 | Weakly |
| Lapatinib | C29H26ClFN4O4S | 581.06 | 2 | 9 | 0 | 5.38 | -8.30 | 101.38 | 4.43 +/- 0.41 | Yes |
| Oxytetracycline | C22H24N2O9 | 460.44 | 7 | 10 | 0 | -2.20 | -1.62 | 201.85 | 4.91 +/- 0.21 | Yes |

\* Properties computed by ChemDraw 23.1.2.
\*\* $F_{in}/F_{out}$ represents the partitioning ratio; mean and standard error are calculated from > 10 condensates. MW: Molecular Weight; HBD: H-bond Donor; HBA: H-bond Acceptor; FC: Formal Charge; tPSA: topological Polar Surface Area.

### General information

The structures of the compounds were established by NMR (^1^H, ^13^C, ^19^F, ^11^B) spectroscopy. The NMR spectra were acquired on a Bruker Avance 300 amd 500 MHz spectrometers at room temperature; the chemical shifts δ were measured in ppm relative to the solvent signals (^1^Н: CDCl_3_, δ = 7.27 ppm, ^13^C: CDCl_3_, δ = 77.00 ppm). Splitting patterns are designated as s, singlet; d, doublet; t, triplet; q, quartet; m, multiplet. The coupling constants (*J*) are in Hertz. HPLC-MS analyses were performed on a Thermo-Dionex Ultimate 3000 instrument (pump + autosampler at 20 °C + column oven at 25 °C) equipped with a diode array detector (Thermo-Dionex DAD 3000-RS) and a MSQ Plus single quadrupole mass spectrometer. Accurate high-resolution mass spectra were obtained on Bruker microTOF-QTM ESI-TOF (Electrospray Ionization/Time of Flight) and Thermo Scientific* LTQ Orbitrap mass spectrometers. Analytical thin-layer chromatography (TLC) was carried out on silica gel plates (silica gel 60 F254 aluminum-supported plates); the visualization was accomplished with an UV lamp (365 nm). Column chromatography was performed on silica gel 60 (230-400 mesh). The measurements were performed in air atmosphere in acetonitrile. All chemicals and reagents were obtained from commercial sources and used as-received, unless otherwise noted. Solvents were purified using standard methods.[1] 8-Chloromethyl-2,6-diethyl-4,4-difluoro-1,3,5,7-tetramethyl-4-bora-3a,4a-diaza-s-indacene was prepared according to a published procedure.[2]

### Synthesis of 4,4-difluoro-8-[4-(di(2-hydroxyethyl))aminophenyl]-2,6-diethyl-1,3,5,7-tetramethyl-4-bora-3a,4a-diaza-s-indacene (MK239)

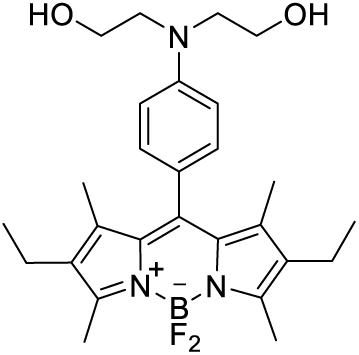

A solution of LiOH (5 mg, 0.21 mmol, 2.1 equiv) in water (0.4 mL) was added to a solution of 4,4-difluoro-8-[4-(di(2-acetoxyethyl))aminophenyl]-2,6-diethyl-1,3,5,7-tetramethyl-4-bora-3a,4a-diaza-s-indacene (57 mg, 0.10 mmol, 1.0 equiv) in THF/MeOH (0.4:1.6 mL) under argon atmosphere. The resulting mixture was stirred at room temperature for 12 h and the solvent was removed under reduced pressure. The residue was twice purified by column chromatography (CH_2_Cl_2_ to CH_2_Cl_2_/ MeOH = 100:1) to get a red solid (40 mg, 82 % yield) with mp 182–184 °C, R*_f_* = 0.20 (CH_2_Cl_2_/MeOH = 50: 1). ^1^H NMR (300 MHz, CDCl_3_): δ 7.08 (d, *J* = 8.6 Hz, 2H, Ar), 6.78 (d, *J* = 8.6 Hz, 2H, Ar), 3.93 (t, *J* = 4.8 Hz, 4H, 2 × CH_2_), 3.67 (t, *J* = 4.8 Hz, 4H, 2 × CH_2_), 3.53 (br. s, 2H, 2 × OH), 2.54 (s, 6H, 2 × CH_3_), 2.32 (q, *J* = 7.4 Hz, 4H, 2 × CH_2_), 1.41 (s, 6H, 2 × CH_3_), 1.00 (t, *J* = 7.4 Hz, 6H, 2 × CH_3_). ^13^C NMR (75 MHz, CDCl_3_): δ 153.4, 132.6, 131.5, 129.6, 129.5, 129.4, 120.4, 113.4, 68.1, 60.6, 55.9, 51.4, 29.8, 17.2, 14.7, 12.5, 12.1. ^19^F NMR (282 MHz, CDCl_3_): δ -146.5 (q, *J =* 33.2 Hz). ^11^B NMR (96 MHz, CDCl_3_): δ 0.83 (t, *J* = 33.2 Hz). HRMS: calcd. [M + H]^+^ for C_27_H_37_BF_2_N_3_O_2_, 484.2946; found 484.2944.

### Synthesis of 4,4-difluoro-8-[4-(di(2-acetoxyethyl))aminophenyl]-2,6-diethyl-1,3,5,7-tetramethyl-4-bora-3a,4a-diaza-s-indacene (DeUv024k2)

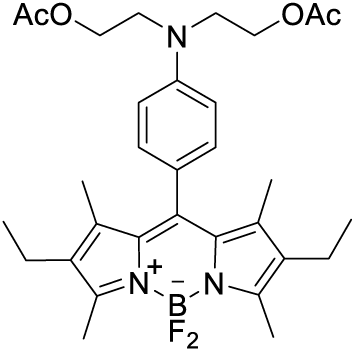

4-(Di(2-acetoxyethyl))amino)benzaldehyde (500 mg, 1.71 mmol, 1.0 equiv) and 3-ethyl-2,4-dimethyl-1*H*-pyrrole (0.49 mL, 3.59 mmol, 2.1 equiv) were dissolved in anhydrous CH_2_Cl_2_ (53 mL) under argon atmosphere. One drop (5 μL) of trifluoroacetic acid was added, and the solution was stirred at room temperature overnight. A solution of 2,3-dichloro-5,6-dicyano-1,4-benzoquinone (427 mg, 1.88 mmol, 1.1 equiv) in anhydrous CH_2_Cl_2_ (74 mL) was added, and the stirring was continued for another 5 h. After the addition of triethylamine (10.3 mL), BF_3_**·**OEt_2_ (10.3 mL) was gradually added during 30 min in an ice-water bath followed by continuous stirring at room temperature overnight. The reaction solution was shaken with 5% aqueous sodium bicarbonate (150 mL), and the mixture was passed through a Celite pad and washed with CH_2_Cl_2_ (80 mL). The organic layer was washed with water (3 × 30 mL), passed through a Celite pad, dried with anhydrous sodium sulfate, and concentrated in vacuo. The residue was twice purified by flash chromatography (CH_2_Cl_2_ to CH_2_Cl_2_/MeOH = 100:1) to obtain a red solid (417 mg, 43% yield) with mp 136–138 °C, R*_f_* = 0.83 (CH_2_Cl_2_/MeOH = 50: 1). ^1^H NMR (300 MHz, CDCl_3_) δ 7.09 (d, *J* = 8.6 Hz, 2H, Ar), 6.85 (d, *J* = 8.6 Hz, 2H, Ar), 4.31 (t, *J* = 6.2 Hz, 4H, 2 × CH_2_), 3.70 (t, *J* = 6.2 Hz, 4H, 2 × CH_2_), 2.54 (s, 6H, 2 × CH_3_), 2.32 (q, *J* = 7.4 Hz, 4H, 2 × CH_2_), 2.09 (s, 6H, 2 × CH_3_), 1.40 (s, 6H, 2 × CH_3_), 1.00 (t, *J* = 7.5 Hz, 6H, 2 × CH_3_). ^13^C NMR (75 MHz, CDCl_3_): δ 171.0, 153.3, 147.4, 138.5, 132.6, 131.5, 129.6, 129.4, 124.4, 112.5, 112.4, 61.2, 53.5, 49.9, 21.0, 17.2, 14.7, 12.5, 12.1. ^19^F NMR (282 MHz, CDCl_3_): δ -146.6 (q, *J* = 33.2 Hz).^11^B NMR (96 MHz, CDCl_3_): δ 0.83 (t, *J* = 33.2 Hz). HRMS: calcd. [M + H]^+^ for C_31_H_41_BF_2_N_3_O_4_, 568.3158; found 568.3147.

### Synthesis of (((2,8-diethyl-5,5-difluoro-1,3,7,9-tetramethyl-5H-4λ^4^,5λ^4^-dipyrrolo[1,2-c:2’,1’-f][1,3,2]diaza-borinin-10-yl)methyl)azanediyl)bis(ethane-2,1-diyl) diacetate (MaKo068).[3]

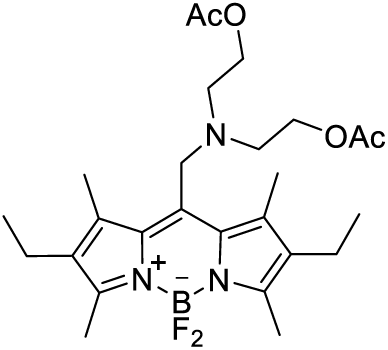

To a solution of 8-chloromethyl-2,6-diethyl-4,4-difluoro-1,3,5,7-tetramethyl-4-bora-3a,4a-diaza-s-indacene (172 mg, 0.49 mmol, 1.0 equiv) in dry MeCN (50 mL) at rt KI (81 mg, 0.49 mmol, 1.0 equiv), K_2_CO_3_ (202 mg, 1.46 mmol, 3.0 equiv), and 2,2’-iminobis(ethyl acetate) hydrochloride (220 mg, 0.97 mmol, 2.0 equiv) were added. The mixture was refluxed for 3 h until the complete conversion of starting BODIPY (TLC monitoring). The resulted mixture was cooled to room temperature, solvent was removed under reduced pressure. The product was isolated by column chromatography (silica gel, eluent petr. ether / EtOAc, 1: 1) afforded product as orange-red solid (114 mg, 46%) with mp 135 - 137 ^о^С, R*_f_* = 0.14 (petr. ether / EtOAc, 5: 1). ^1^H NMR (CDCl_3_, 300 MHz): δ 4.12 (t, *J* = 5.2 Hz, 4H, 2 × CH_2_), 4.08 (s, 2H, CH_2_), 3.01 (t, *J* = 5.2 Hz, 4H, 2 × CH_2_), 2.52 (s, 6H, 2 × СН_3_), 2.47 – 2.30 (q, *J* = 7.3 Hz, 4H, 2 × СН_2_), 2.38 (s, 6H, 2 × СН_3_), 1.96 (s, 6H, 2 × CH_3_), 1.07 (t, *J* = 7.3 Hz, 6H, 2 × СН_3_). ^13^C NMR (CDCl_3_, 75 MHz): δ 170.9 (2 × CО), 153.7, 137.1, 133.2, 132.8, 62.3, 51.2, 50.2, 20.8, 17.3, 14.7, 13.7, 12.6. ^19^F NMR (CDCl_3_, 282 MHz): δ -146.5 (q, *J* = 33.1 Hz). ^11^B NMR (CDCl_3_, 96 MHz): δ 0.59 (t, *J* = 33.1 Hz). HRMS: calcd [M+H]^+^ for С_26_H_39_BF_2_N_3_O ^+^, 506.3001; found, 506.3004.

## NMR Spectra

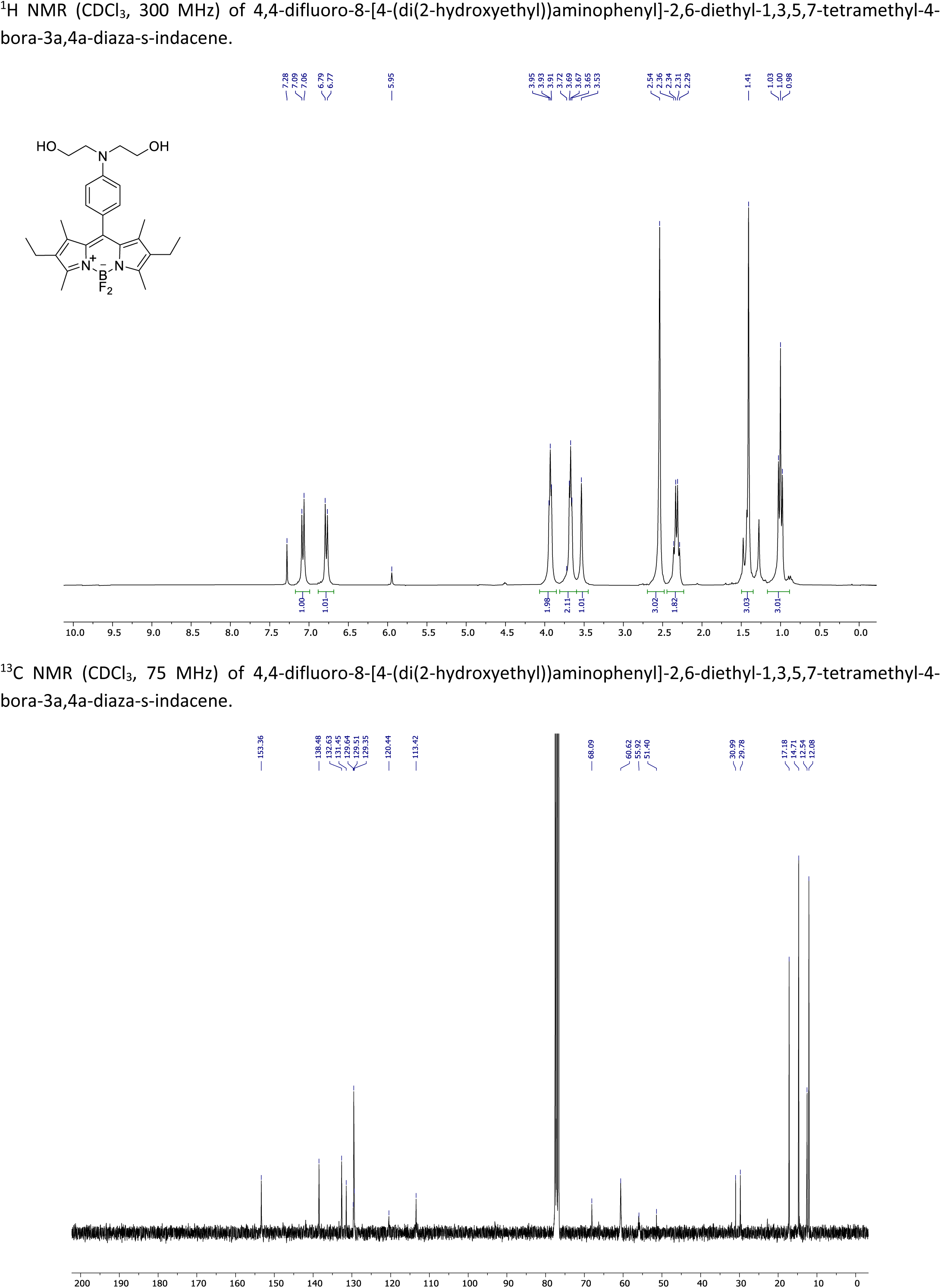

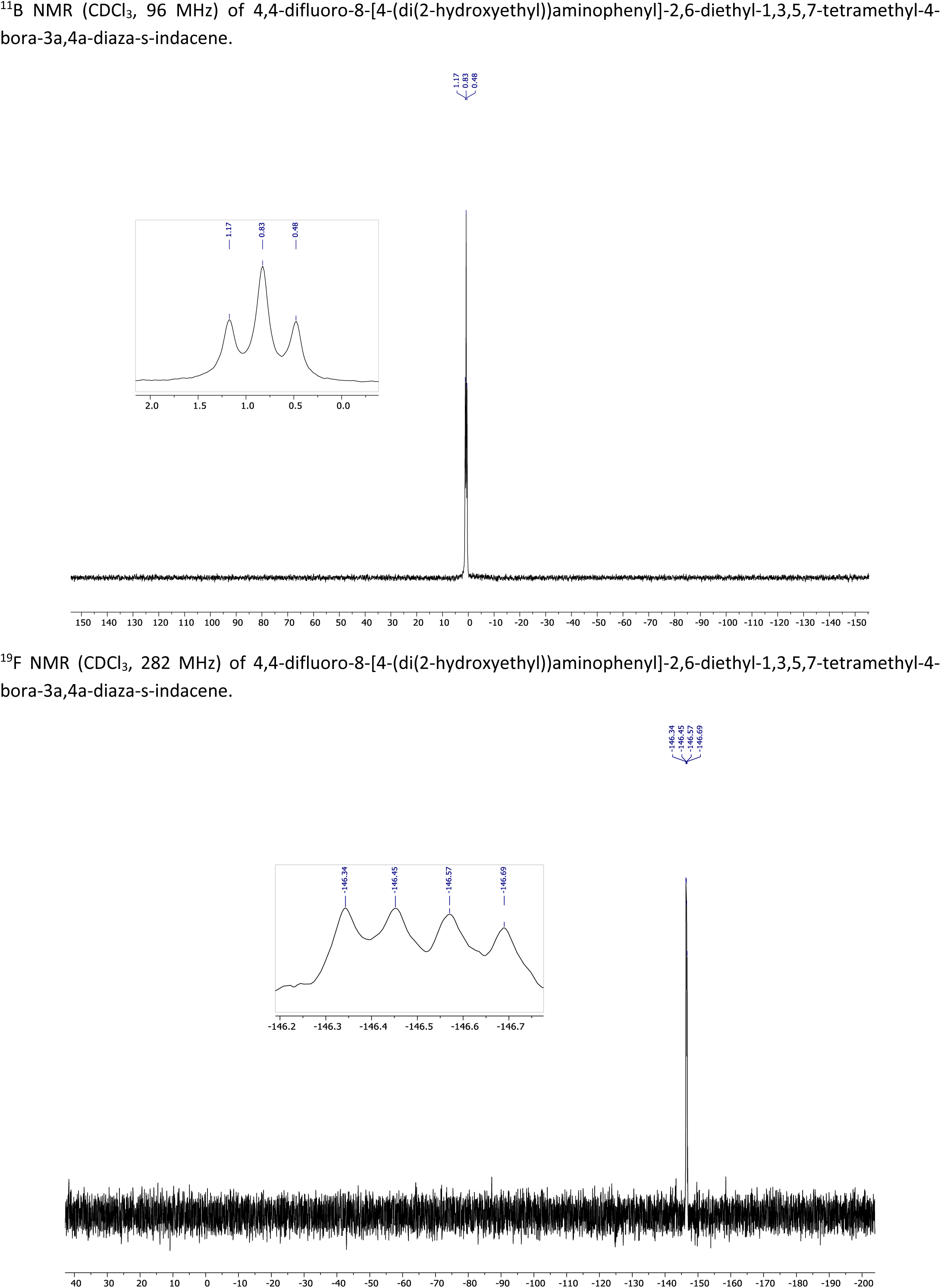

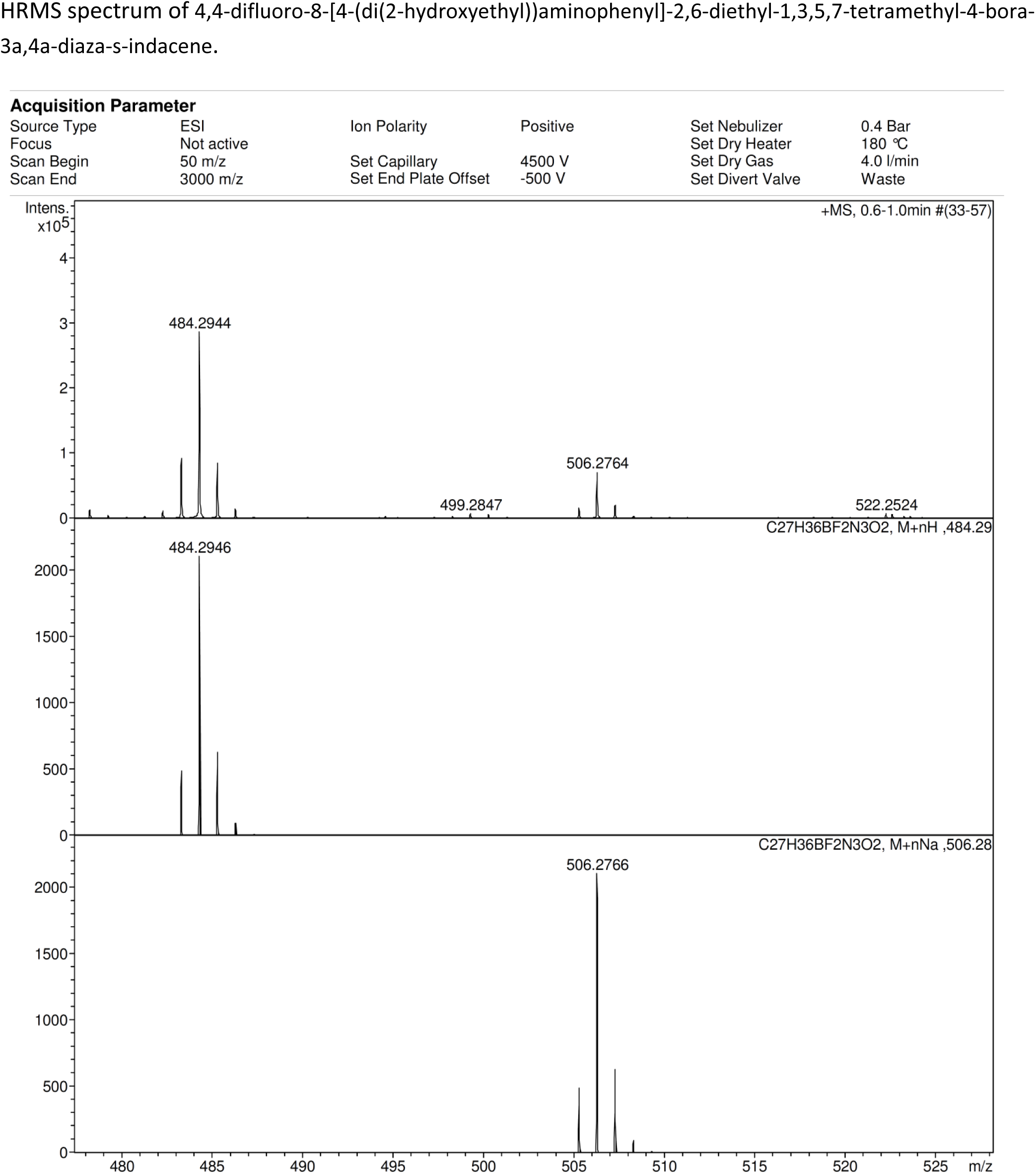

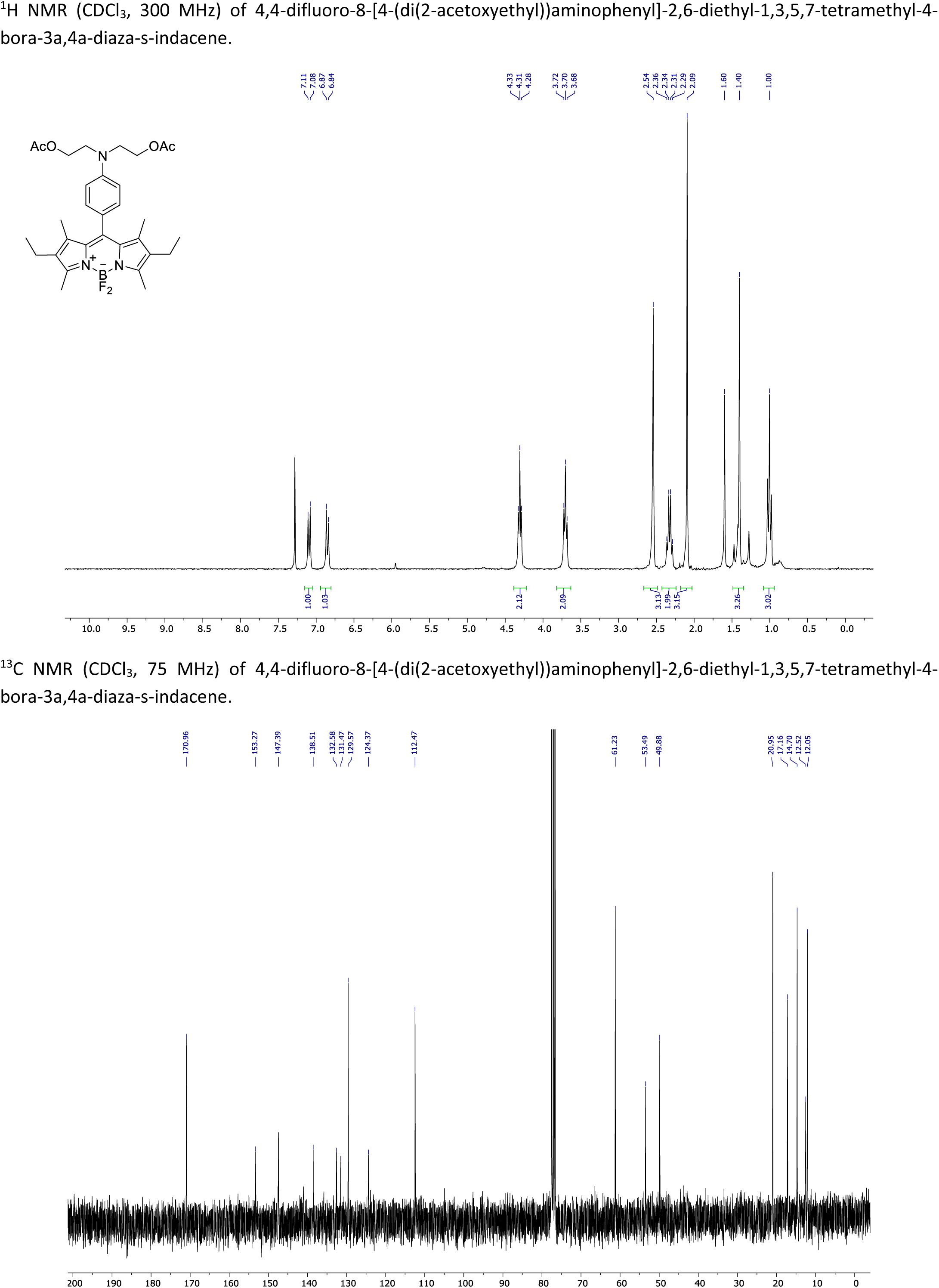

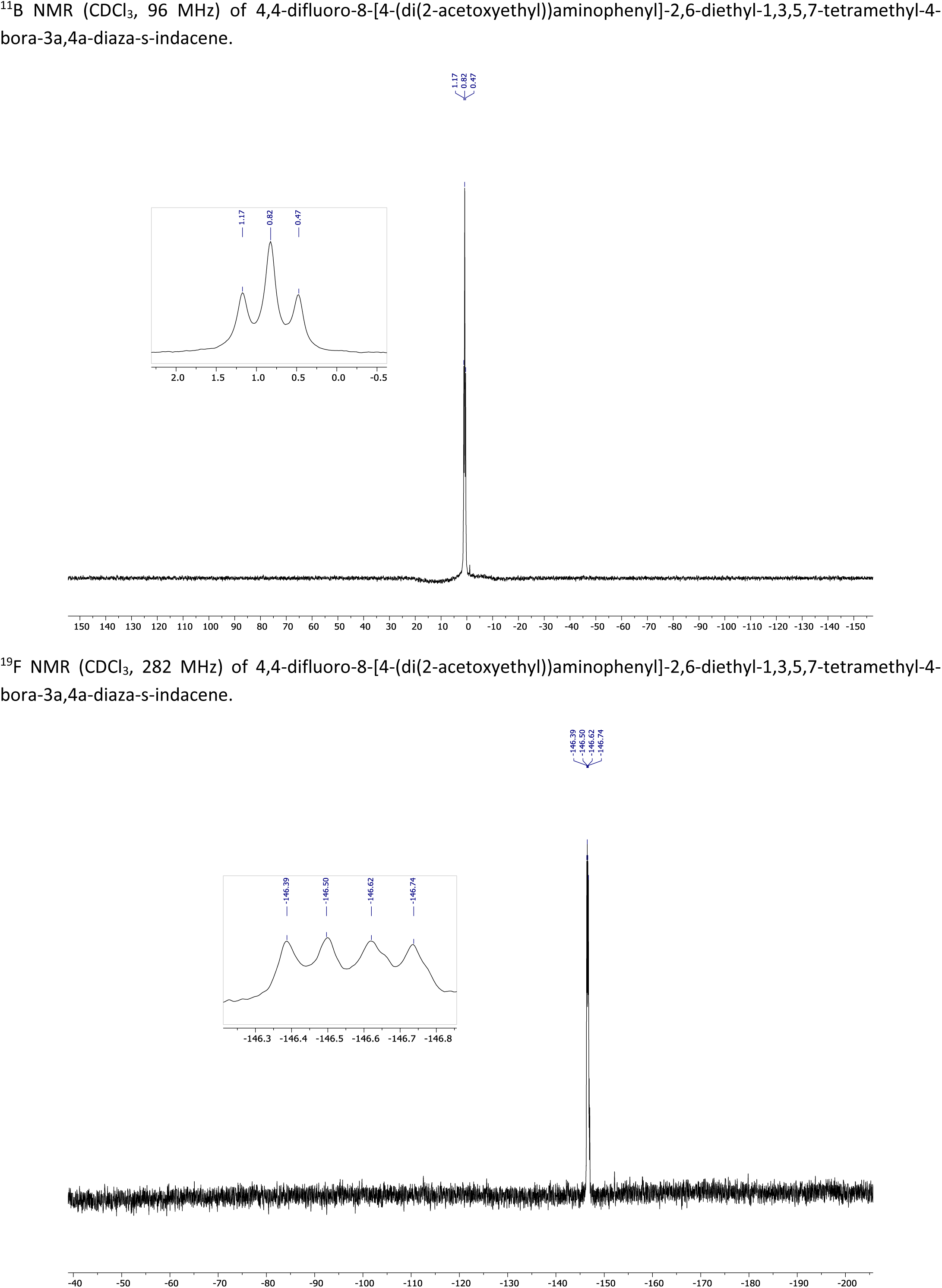

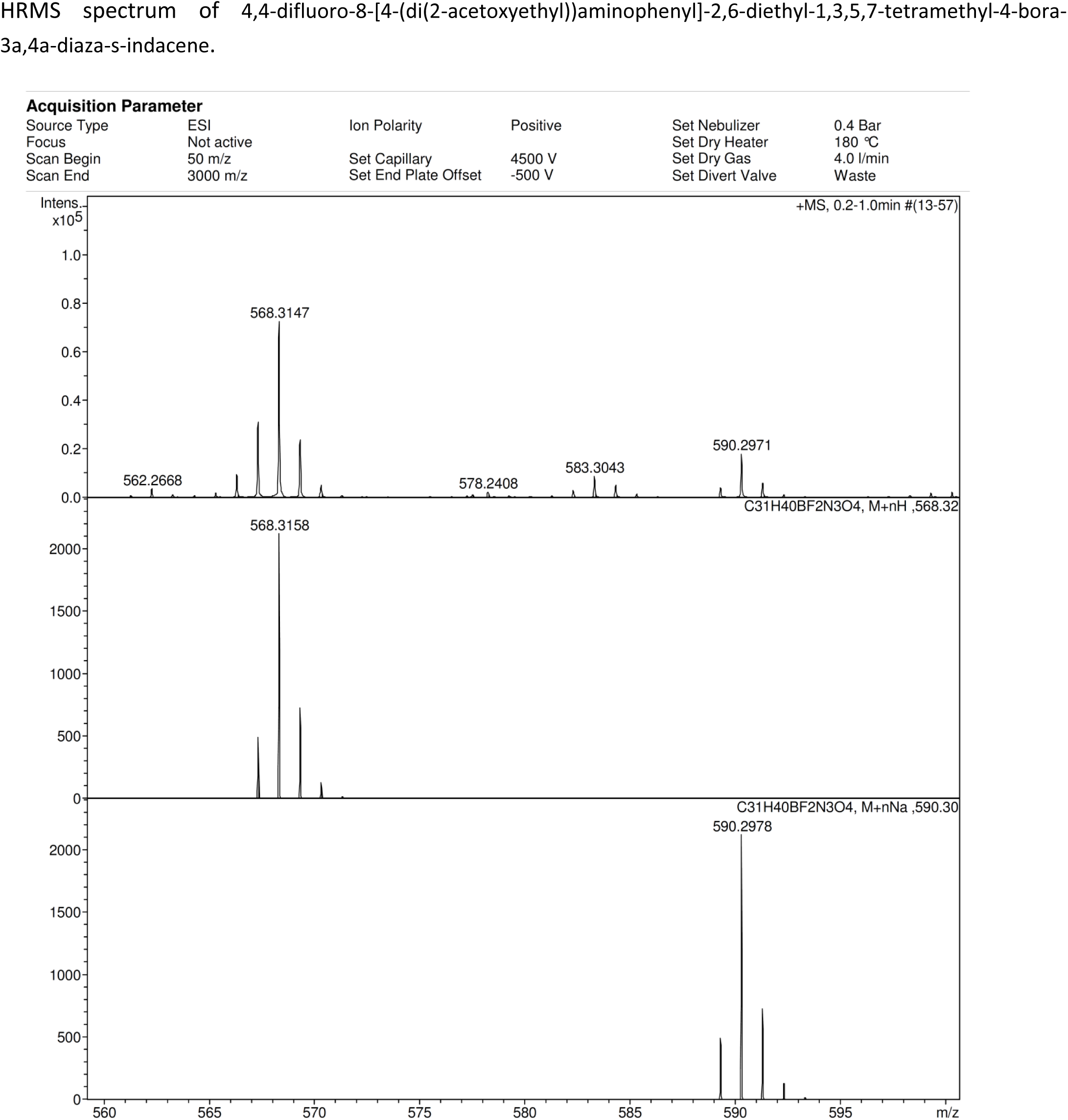

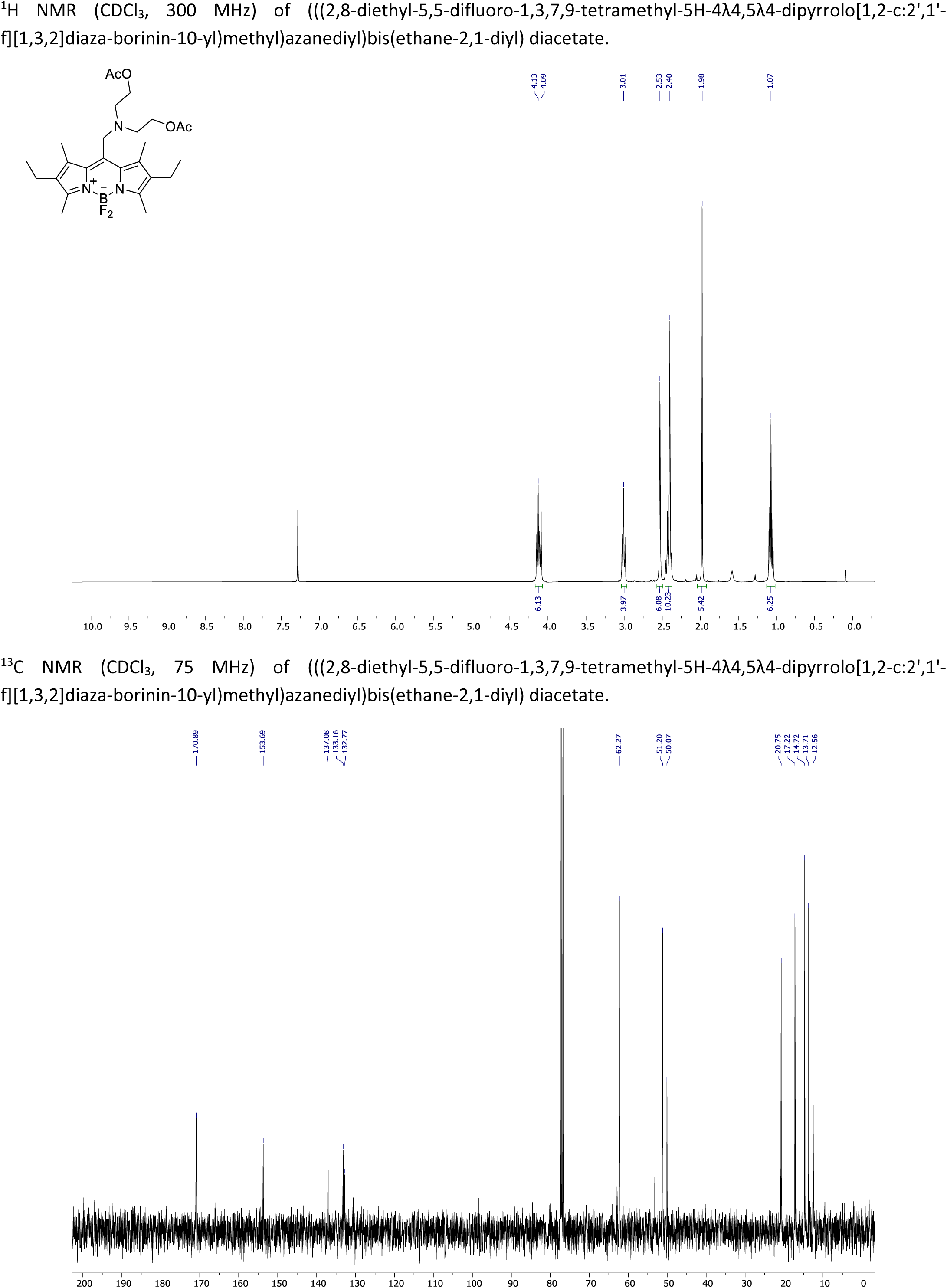

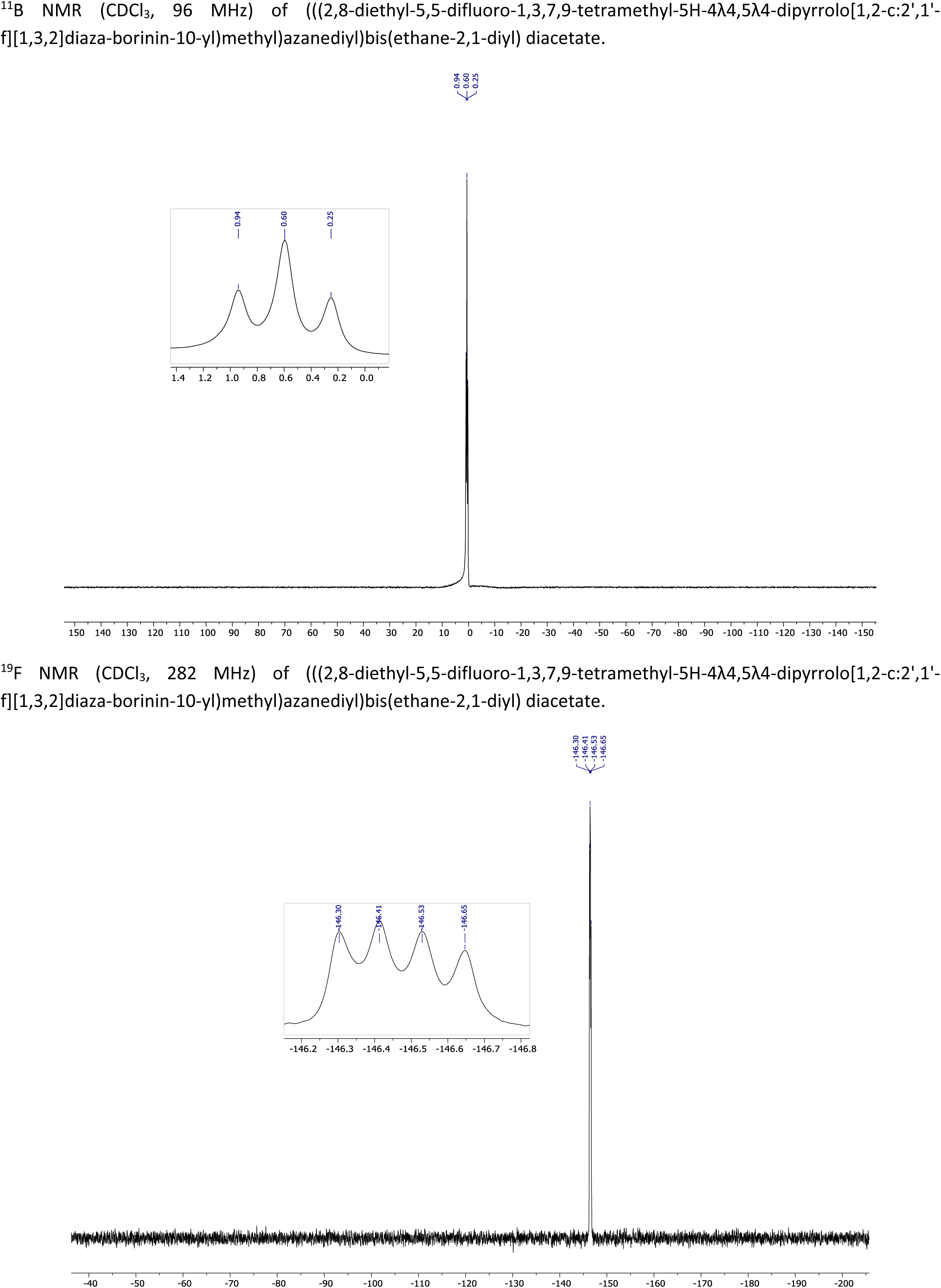

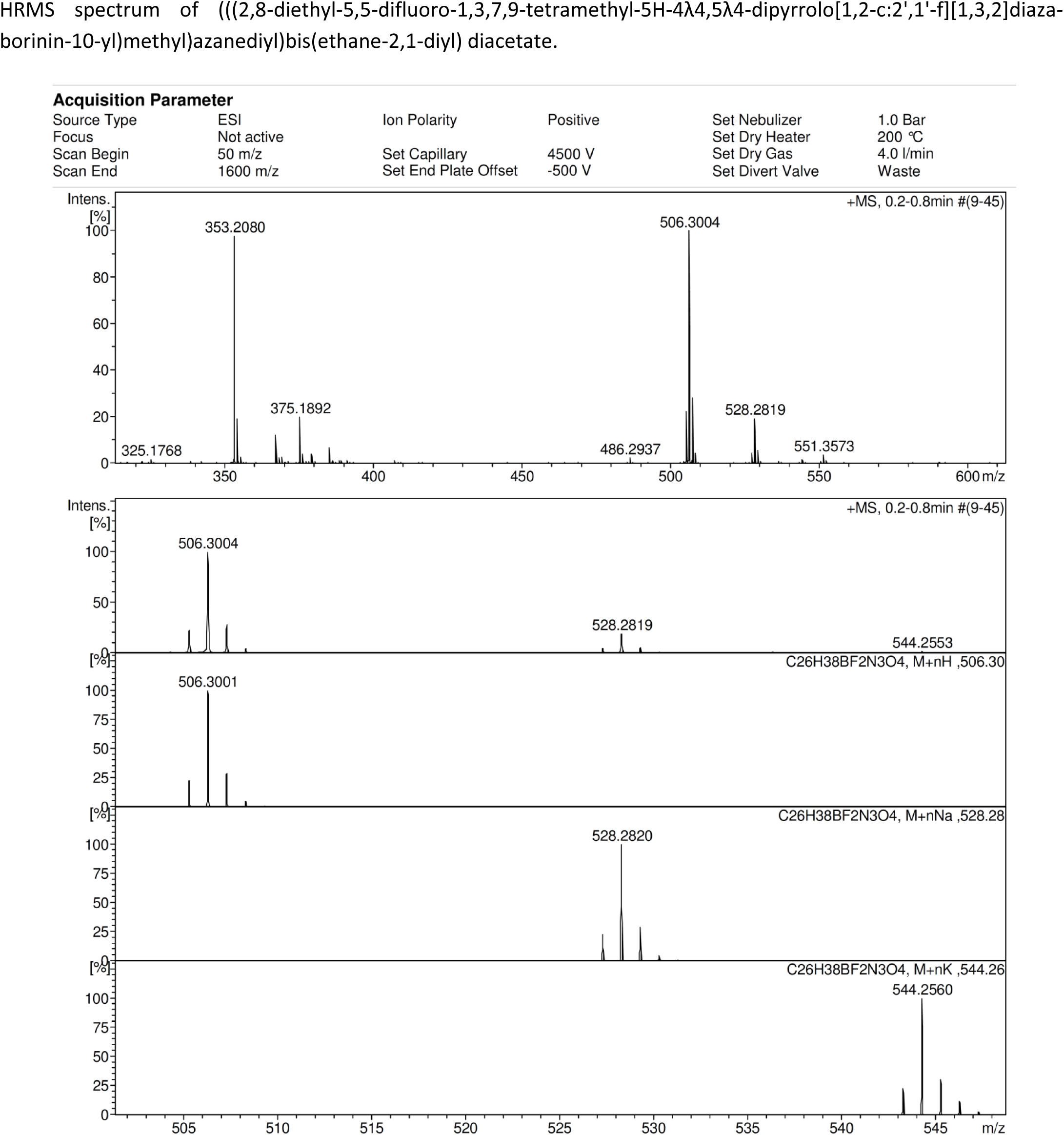

